# BoneGraph: A Domain-Specialised, Self-Correcting Reasoning System for Bone Science Retrieval, Grounded Inference, and Image Mechanics

**DOI:** 10.64898/2026.09.17.752482

**Authors:** Jon Valijonov, Peter Soar, James Le Houx, Gianluca Tozzi

## Abstract

Bone science literature spans biology, mechanics, materials science, and clinical medicine, and its volume makes reliable knowledge synthesis increasingly difficult. General-purpose large language models (LLMs) answer fluently but under-represent this niche domain, cannot cite specific evidence, and offer no mechanism to be corrected durably. Here we present BoneGraph, a domain-specialised system for bone science delivered as a five-tab web application over a shared substrate: a curated full-text corpus of 7,449 documents embedded into 248,629 passage vectors using SPECTER2, a scientific-paper embedding model, and a bone knowledge graph of 1,597 concepts with 1,699 causal relations. The five tabs are: (I) Chat, retrieval-augmented question answering with server-rebuilt inline citations; (II) Search, raw semantic retrieval with no LLM in the loop; (III) Reasoning, a self-correcting loop in which a deterministic physics check and a literature/knowledge-graph critic constrain the answer, and a user’s feedback becomes a durable, per-user rule; (IV) Vision, a bone-region classifier trained on frozen BiomedCLIP features that grounds a vision-language model, guarded against out-of-distribution inputs and augmented with image-embedding correction memory; and (V) Mechanics, integrating our previous data-driven image mechanics (D^2^IM) model that predicts displacement and strain fields from a single undeformed micro-CT image. All inference is performed locally, without third-party API calls, and the public beta is served at bonegraph.org. Retrieval attains a mean reciprocal rank (MRR) of 0.928 on a 30-question, seven-domain benchmark, and the Vision classifier attains 92.6% accuracy on the held-out MURA (MUsculoskeletal RAdiographs) dataset. A grounded-reasoning benchmark shows that, with the correct passage, BoneGraph raises answer accuracy from 42% to 78%. BoneGraph makes a major contribution to bone-science informatics: to our knowledge it is the first domain-specialised system to unify curated retrieval, deterministic physics-grounded self-correction, and durable per-user learning for bone science.

## 1. Introduction

Bone science sits at the intersection of biology, mechanics, materials science, and clinical medicine. Research in the field spans bone morphology, structure-function relationships, biomechanical properties, pathology (e.g. osteoporosis, stress fracture), medical imaging, biomaterials, and multiscale computational simulation. Because relevant knowledge is distributed across thousands of journal publications, comprehensive literature synthesis is a substantial and growing burden for researchers.

Large language models (LLMs), built on the transformer architecture [1], have demonstrated strong general-purpose question answering and text generation [2]. However, their application to a highly specialised scientific domain such as bone science is limited by three factors. First, standard LLMs retrieve from parametric memory, which cannot be updated without retraining and provides no grounding to specific peer-reviewed evidence. Second, general training corpora under-represent niche domains relative to broadly documented fields, so this parametric knowledge is thin and error-prone exactly where precision matters most. Third, when an expert user identifies an error, a stock LLM has no durable mechanism by which to accept the correction, and the same mistake may recur on the next similar question. Retrieval-Augmented Generation (RAG) [3] addresses the first of these limitations by grounding responses in a retrieved corpus at inference time, but the quality of a RAG system is bounded by the coverage of its corpus and by the representational power of its embedding model, and retrieval alone does not verify that an answer is physically or logically consistent. Existing academic search tools, moreover, are not curated for bone science and interleave unrelated biomedical literature.

Existing work addresses the remaining two limitations only in part. Biomedical language models, including domain-specialised encoders such as BioBERT [4], PubMedBERT [5], and Med-PaLM [6], and reasoning models such as HuatuoGPT-o1-8B [7], improve clinical coverage but under-represent bone biomechanics, while inference-time techniques such as chain-of-thought prompting [8], self-consistency [9], and self-refinement [10] sharpen reasoning without grounding it in evidence. On the retrieval side, open scholarly indexes such as OpenAlex [11] and PubMed [12] establish the limits of corpus coverage, while citation-informed scientific embeddings such as SPECTER [13] and SPECTER2 [14] determine representational quality, but neither is curated for bone science.

Structured and physical knowledge can constrain reasoning further. Knowledge graphs have long encoded causal and relational structure for biomedical reasoning [15], though none spans the full range of bone morphology, mechanics, and pathology, and well-established bone-mechanics relationships are deterministic enough to check directly. For imaging, vision-language models built on contrastive and Vision-Transformer encoders [16–17], from LLaVA [18] and LLaVA-Med [19] to the pathology copilot PathChat [20], which adapts a self-supervised foundation encoder [21–22], and outperforms general assistants such as GPT-4V [23], add visual capability, yet the strongest among them require large-curated instruction sets and substantial GPU training. Deep-learning surrogates for biomechanical fields offer fast alternatives to finite-element modelling and to digital volume correlation [24], but remain isolated from any retrieval or reasoning capability. No existing system unifies curated bone-science retrieval, verifiable self-correcting reasoning, and durable user correction in a single, locally deployable package.

Here we present BoneGraph, a domain-specialised, self-correcting reasoning system for bone science, deployed as a public beta at bonegraph.org. Rather than a single chatbot, BoneGraph is a family of five task-specific pipelines over one shared substrate (a curated corpus, a dense embedding index, a bone knowledge graph, and a set of local open-weight models) unified behind a single web application and a per-user account layer. The design goal is a system that can be audited: it grounds claims in cited evidence, checks them against deterministic physics, and learns from the user in a way that is transparent and reversible. The principal contributions of this paper are threefold, concerning in turn the curated substrate upon which the system rests, the task-specific pipelines constructed over it, and the demonstration that the resulting system is locally deployable and designed to be domain-transferable.

BoneGraph is built on a curated bone-science corpus comprising the metadata of 54,634 peer-reviewed papers together with 16 open-access textbooks, assembled via the OpenAlex API [11], together with a three-tier PDF resolution pipeline that recovers 7,674 validated full-text PDFs beyond what the API directly provides. A text-processing and retrieval pipeline comprising full-text extraction, two-pass language filtering, sentence-aware chunking, and SPECTER2 [14] dense embedding produces 248,629 semantically indexed passages.

On this substrate, the self-correcting Reasoning pipeline combines an agent that reasons only from the question and the user’s own rules; a deterministic eight-rule physical-grounding check; and a critic armed with retrieved literature and one-hop knowledge-graph edges that can accept, dispute, or flag conflicting evidence. User feedback is compiled into durable per-user rules applied on every future request. The Vision pipeline is built on a hybrid, lightweight-grounding design: a bone-region classifier trained on frozen BiomedCLIP [25] features from MURA [26] grounds a vision-language model; an out-of-distribution guard withholds that prediction on scans outside the training scope; and image-embedding correction memory recalls a prior correction so a re-windowed or rotated copy of a previously corrected scan still triggers the fix. The Mechanics pipeline integrates our D^2^IM [27], a deep-learning model that predicts displacement and strain fields from a single undeformed micro-CT image with no finite-element model or digital volume correlation at inference. The whole system runs entirely locally and at low cost: all models are served on a single NVIDIA Jetson AGX Orin through Ollama [28] and Hugging Face weights, exposed via a Cloudflare Tunnel with no third-party API calls at inference time. Finally, the architecture is domain-transferable: the same pipeline carries over to another specialised domain unchanged, adapted only by substituting the domain-specific keywords that define the corpus and the specialised models behind their single swap points. The entire system was, moreover, assembled and operated on commodity hardware.

## 2. Results and Discussion

We evaluated retrieval on a manually constructed benchmark including thirty questions spanning seven bone-science domains (mechanics, morphology, pathology, biomaterials, simulation, imaging, and mechanobiology), each paired with expected-answer keywords drawn from the ground-truth literature. At k = 10, the system attains an overall Mean Reciprocal Rank (MRR) of 0.928. Every question not returned at rank 1 had a relevant passage at rank 2 or 3, indicating minor ranking errors rather than gaps in coverage. Per-domain and overall MRR are reported in Table 1. The lowest scores arise in simulation (0.833) and biomaterials (0.867), the domains whose terminology is most heavily shared with adjacent engineering literature.

**Table 1.** Mean reciprocal rank (MRR) for retrieval by domain and overall (30 questions, 7 domains, k = 10).

| Domain | MRR |
| --- | --- |
| Imaging | 1.000 |
| Mechanobiology | 1.000 |
| Morphology | 1.000 |
| Mechanics | 0.900 |
| Pathology | 0.900 |
| Biomaterials | 0.867 |
| Simulation | 0.833 |
| **Overall** | **0.928** |

Retrieval quality alone does not establish whether grounding improves answers. We therefore constructed a second benchmark of fifty multiple-choice questions drawn directly from the corpus. Each question requires identification of a numerical quantity in the literature and then used in a single calculation. Questions were authored from corpus passages rather than from model knowledge. Each records its source chunk together with the exact quoted sentence. Whether the deciding quantity is present in the retrieved evidence can therefore be verified automatically.

Four retrieval conditions were compared over identical questions, option orderings, decoding parameters and scoring. All were served by the deployed HuatuoGPT-o1-8B behind the bone-science system prompt at temperature 0. Under no retrieval (closed-book), the model receives the question alone. Under keyword retrieval, a BM25 index supplies the fourteen highest-ranked passages. Under keyword retrieval with reranking, a language model selects four passages from the 150 highest-ranked. Both keyword conditions also receive the one-hop knowledge-graph facts that the deployed critic uses. Under perfect retrieval (oracle), the passage known to contain the answer is supplied directly by identifier, without graph facts. Perfect retrieval is not an achievable system but rather an upper bound on what the same model can attain with this corpus. Each condition was run under both of the answer prompts reproduced in Appendix A. The direct prompt asks only for an answer. The structured prompt requires the model to state each quantity, convert units, calculate, and only then match its result to the options.

Results are reported in Table 2. Under the structured prompt, perfect retrieval answers 39 of 50 questions and no retrieval 21 of 50. This difference of 18 questions, or 36 percentage points, is significant by exact McNemar test. Neither keyword retrieval condition separates from no retrieval. Keyword retrieval reaches 26 of 50 and reranking 29 of 50. The gap between reranking and perfect retrieval remains significant. Reranking does not improve upon keyword retrieval under either prompt. The additional inference call it requires is therefore not presently justified.

**Table 2.**
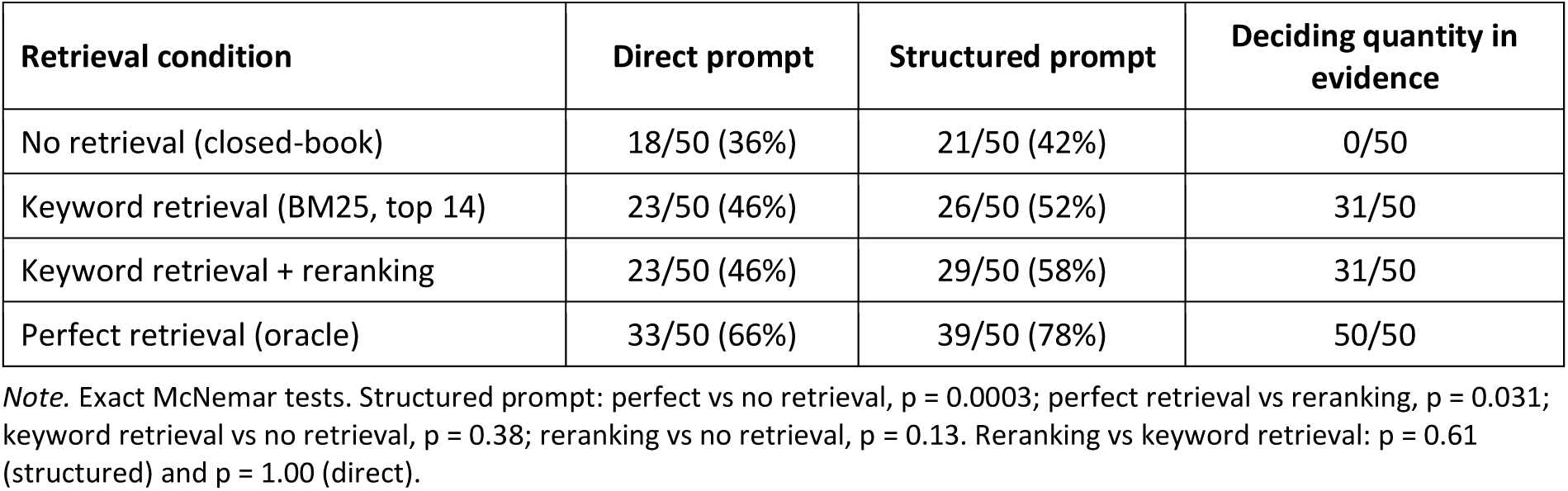
Grounded-reasoning benchmark with questions answered correctly out of 50, by retrieval condition and prompt (Appendix A). All conditions use HuatuoGPT-o1-8B at temperature 0. The two keyword retrieval conditions also receive one-hop knowledge-graph facts. The final column gives the questions whose deciding quantity was in the evidence.

The shortfall against perfect retrieval is one of evidence delivery rather than of reasoning. A per-question diagnostic records whether the deciding quantity appeared in the passages supplied by retrieval, a property we term needle-presence (Table 3). Each keyword condition delivered that quantity for 31 of the 50 questions. For each condition, these were the same questions under both prompts. Accuracy follows needle-presence closely whenever the prompt requires the quantity to be used. Under the structured prompt, reranking answered 24 of the 31 questions whose deciding quantity was present. It answered 5 of the 19 for which it was absent. Keyword retrieval alone answered 20 of 31 against 6 of 19.

**Table 3.**
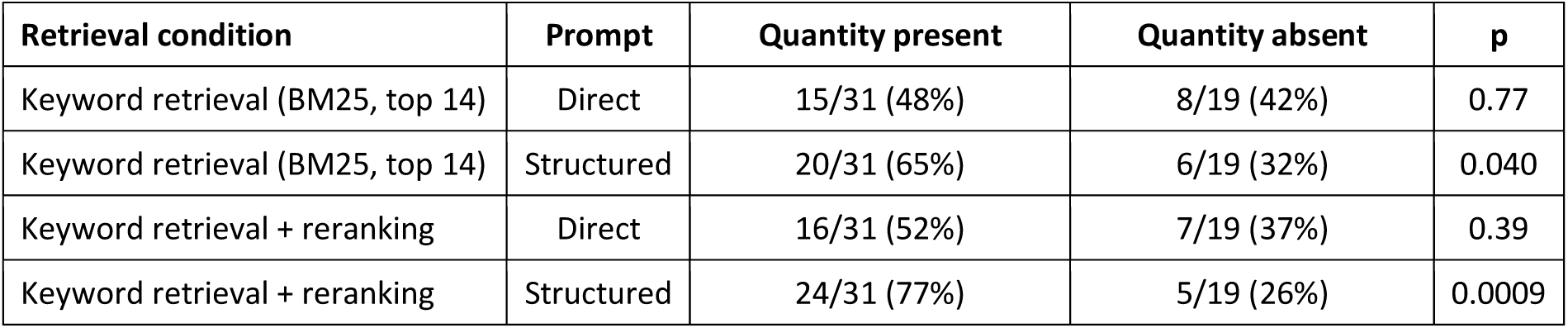
Accuracy of the two keyword retrieval conditions, split by whether the deciding quantity was retrieved (31 of 50 questions under both prompts). p, two-sided Fisher’s exact test.

Under the direct prompt, that correspondence all but disappears. Reranking answered 16 of 31 against 7 of 19, and keyword retrieval 15 of 31 against 8 of 19. The gain the structured prompt delivers accrues almost entirely to questions whose deciding quantity had been retrieved. Questions lacking it do not improve. Neither difference is significant at this sample size. Grounding on this benchmark therefore appears to require two conditions jointly: that the answer-bearing passage arrive, and that the prompt compel its use. The first is met for approximately three questions in five. MRR does not capture this, because it rewards returning passages on the correct topic. A passage may be topically apt and yet contain none of the quantities upon which a question turns.

Prior to their connection to the critic, both evidence channels were audited on five fracture-domain probes. The literature channel proved strong and required no tuning: all five probes returned on-topic passages at high cosine similarity (0.78 to 0.84) and surfaced precisely the papers that a critic would be expected to cite. The knowledge graph proved serviceable, subject to one caveat. One-hop neighbourhoods were excellent and most short concept-to-concept paths were clean, but the graph is fragmented, so that some pairs have no connecting path, a case that is handled gracefully, and composed multi-hop paths can imply falsehoods even when each individual hop is true. This audit motivates the composed-path constraint: the critic receives only raw one-hop edges.

The trained heads of the Vision tab were evaluated on MURA’s held-out validation split, which is disjoint by patient from the training data (Figure 1a). Operating over frozen BiomedCLIP features, body-region classification attains 92.6% accuracy with a one-hidden-layer MLP and 89.6% with a linear probe, at macro-F1 scores of 0.918 and 0.886 respectively. Both figures are well above the seven-class chance rate of 14%, and every region individually exceeds 85%. Abnormality detection is considerably more difficult, as is to be expected on MURA: the MLP head attains 74.2% accuracy (macro-F1 0.735) but recalls only 61% of abnormal studies and is accordingly regarded as preliminary. The out-of-distribution guard separates in-scope from out-of-scope inputs cleanly (Figure 1b). MURA validation radiographs score a nearest-neighbour cosine of 0.83 to 0.99 to the training features (median 0.95), whereas out-of-scope images, such as random noise and flat fields, score 0.47 to 0.50. The first-percentile threshold of 0.827 thus admits legitimate radiographs while withholding a prediction on inputs for which the classifier was never trained. A spine micro-CT is a representative case: in the absence of the guard, it yields a confident but incorrect label of ‘shoulder’.

**Figure 1.**
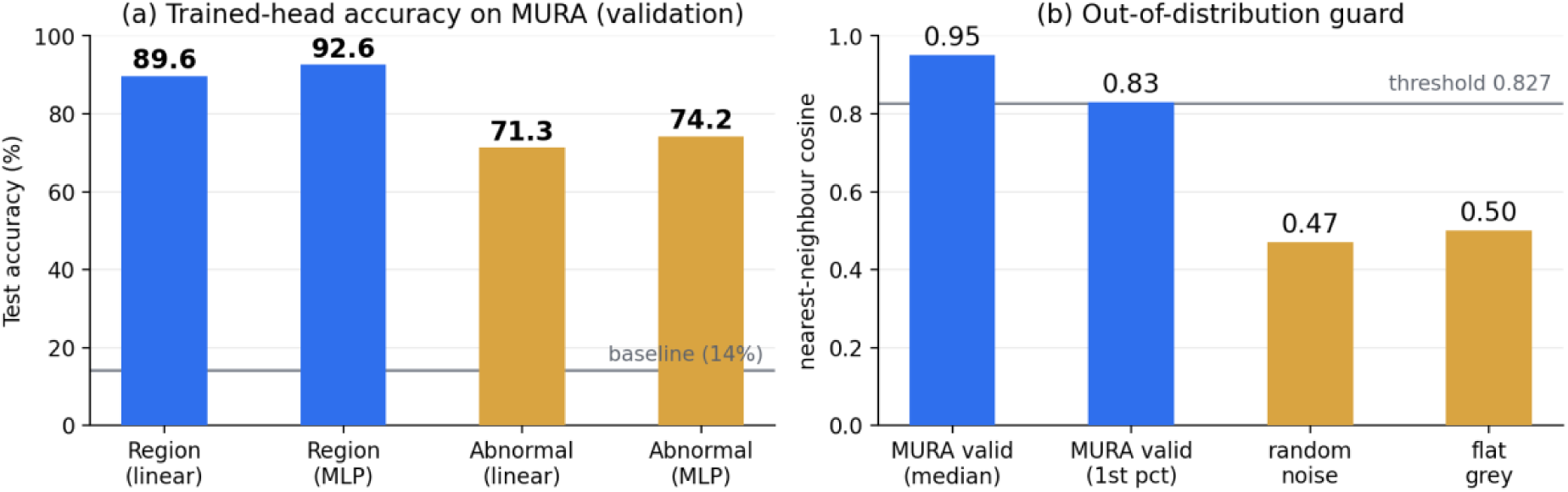
(a) Trained-head accuracy on held-out MURA where body region separates well, while abnormality is substantially harder (MLP abnormal recall 0.61). (b) The out-of-distribution guard where in-scope radiographs sit above the 0.827 nearest-neighbour-cosine threshold, whilst out-of-scope images fall well below and are withheld.

Two caveats bound the weight that should be placed on these figures. First, the scores produced by LLM benchmarks are highly sensitive to prompt wording: minor alterations to an evaluation template can shift accuracy by tens of points, and in opposite directions for different models [29]. The figures reported here are therefore obtained under fixed and disclosed prompts, and are not directly comparable with numbers obtained elsewhere under different prompts. Second, the retrieval benchmark is small, comprising thirty questions, and was executed only once. These results are accordingly presented as indicative rather than definitive, and the construction of a larger, expert-validated benchmark is left to future work.

Complementing the quantitative evaluations, we report a worked example from each of the Chat, Reasoning, Vision, and Mechanics tabs. Figure 2 shows the Chat tab answering a question on what determines the fracture toughness of cortical bone. Each claim in the answer carries an inline citation to a retrieved passage, and the “References” section beneath the answer lists only the three sources cited in the answer text, numbered in ascending order of retrieval rank and linking to the source paper.

**Figure 2.**
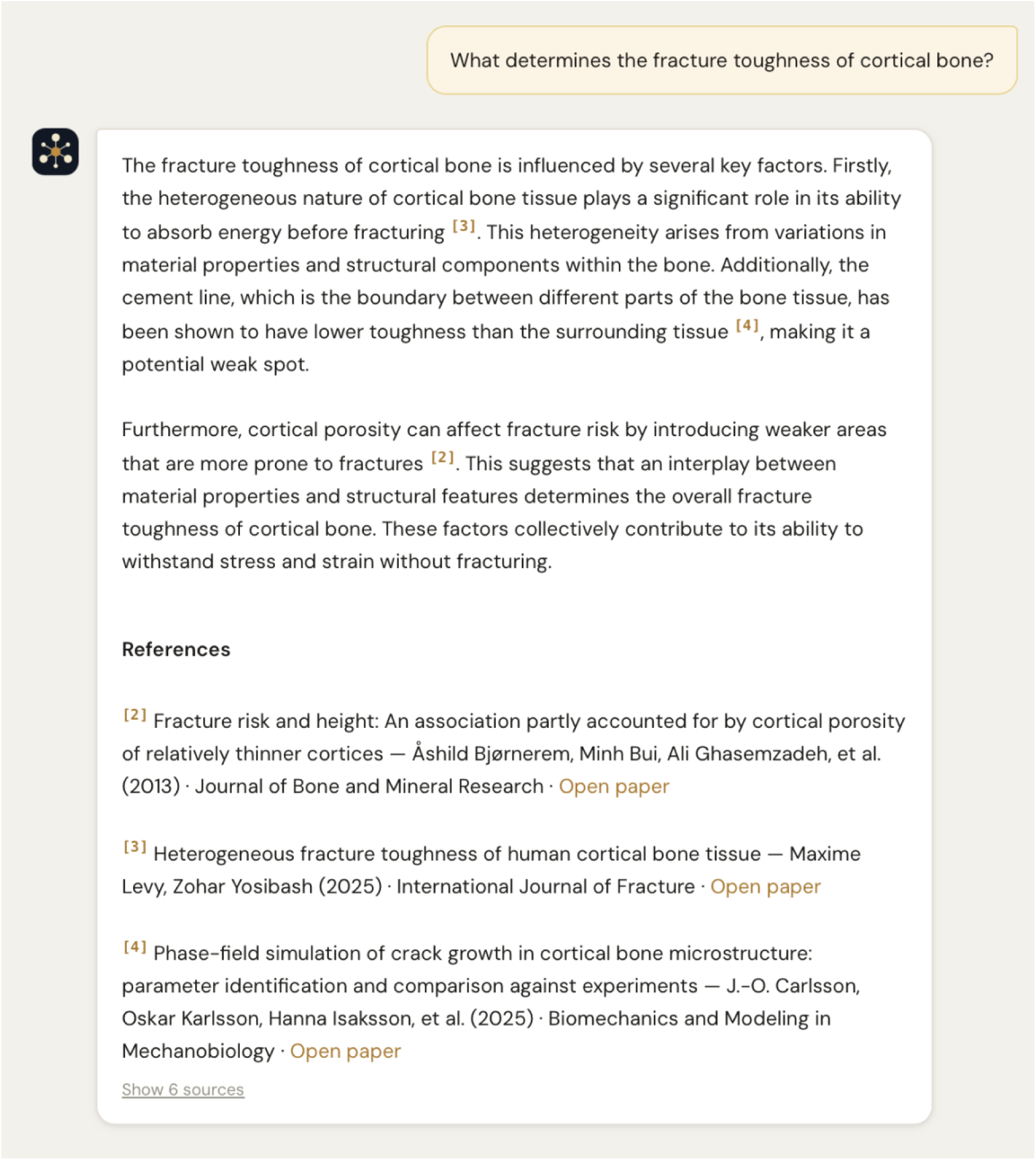
The Chat tab: retrieval-augmented question answering over the curated corpus, with inline [N] citations and a reference.

Figure 3 traces the Reasoning tab’s feedback loop end to end, from an uncorrected erroneous answer to the correct answer enforced by the compiled rule. The question carries a premise, namely that cortical bone fails before trabecular bone in osteoporosis. With no user rule in effect, the agent accepted the premise, the answer passed all eight built-in grounding rules, and the critic accepted it on first review (Figure 3a). The user rejected the answer and described the correction in free text (Figure 3b), which llama3.2:3b compiled into a structured, editable comparative rule that the user then confirmed (Figure 3c). When the same question was asked again with the rule in effect, the answer passed grounding against nine rules, including the user’s, and the critic accepted it after one revision; the answer now states that trabecular bone fails before cortical bone (Figure 3d). The example therefore demonstrates that a single taught correction changes the answer when the same question is asked again, without any change to model weights. It also shows the limit of the unaided loop: on first review, neither the eight built-in rules nor the critic questioned the premise, and the correction originated with the user.

**Figure 3.**
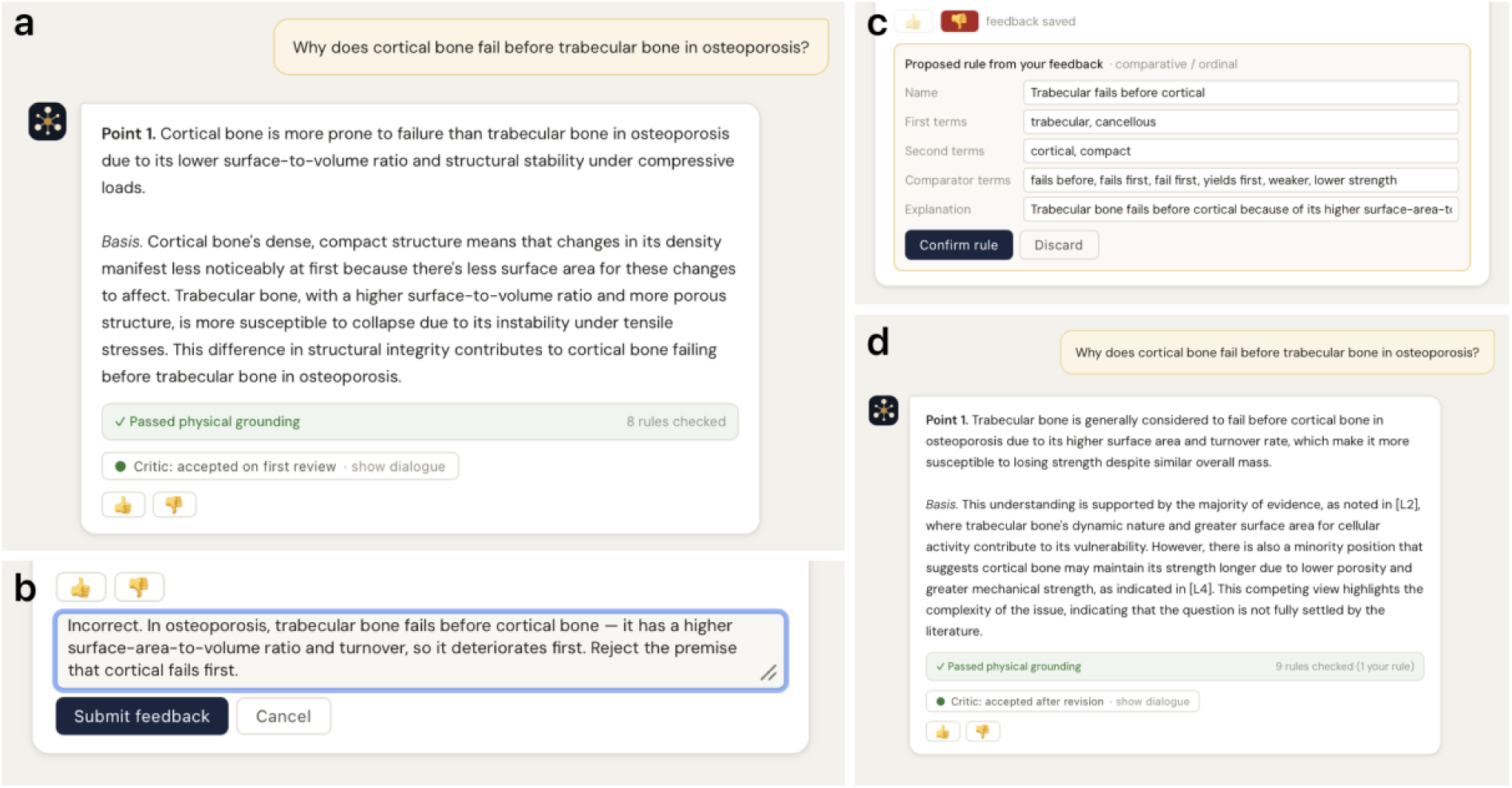
A worked walkthrough of the Reasoning tab’s feedback loop. The question carries a premise: cortical bone fails before trabecular bone in osteoporosis, whereas trabecular bone is often reported to deteriorate first. **(a)** With no user rule in effect, the agent accepts the premise, and the answer passes physical grounding and is accepted by the critic on first review. **(b)** The user thumbs the answer down and describes the correction in free text. **(c)** llama3.2:3b compiles the correction into a structured, editable comparative rule, which the user confirms into their personal rule registry. **(d)** The same question re-asked with the rule in effect: the agent now opens with the correct answer, physical grounding passes with the user, and the critic accepts.

Figure 4 illustrates correction memory recovering from a misidentification and recalling the correction on a rotated copy of the same scan. A femur radiograph, which lies outside the upper-limb scope of the classifier, was labelled humerus by the classifier at 68%, and the vision-language model, given that prediction, also identified the bone as the humerus (Figure 4a). The user rejected the identification and saved a one-line correction. When a copy of the same scan rotated by 20° was uploaded, correction memory matched it to the stored correction at 96% similarity, the classifier hint was suppressed, and the model identified the bone as the femur (Figure 4b).

**Figure 4.**
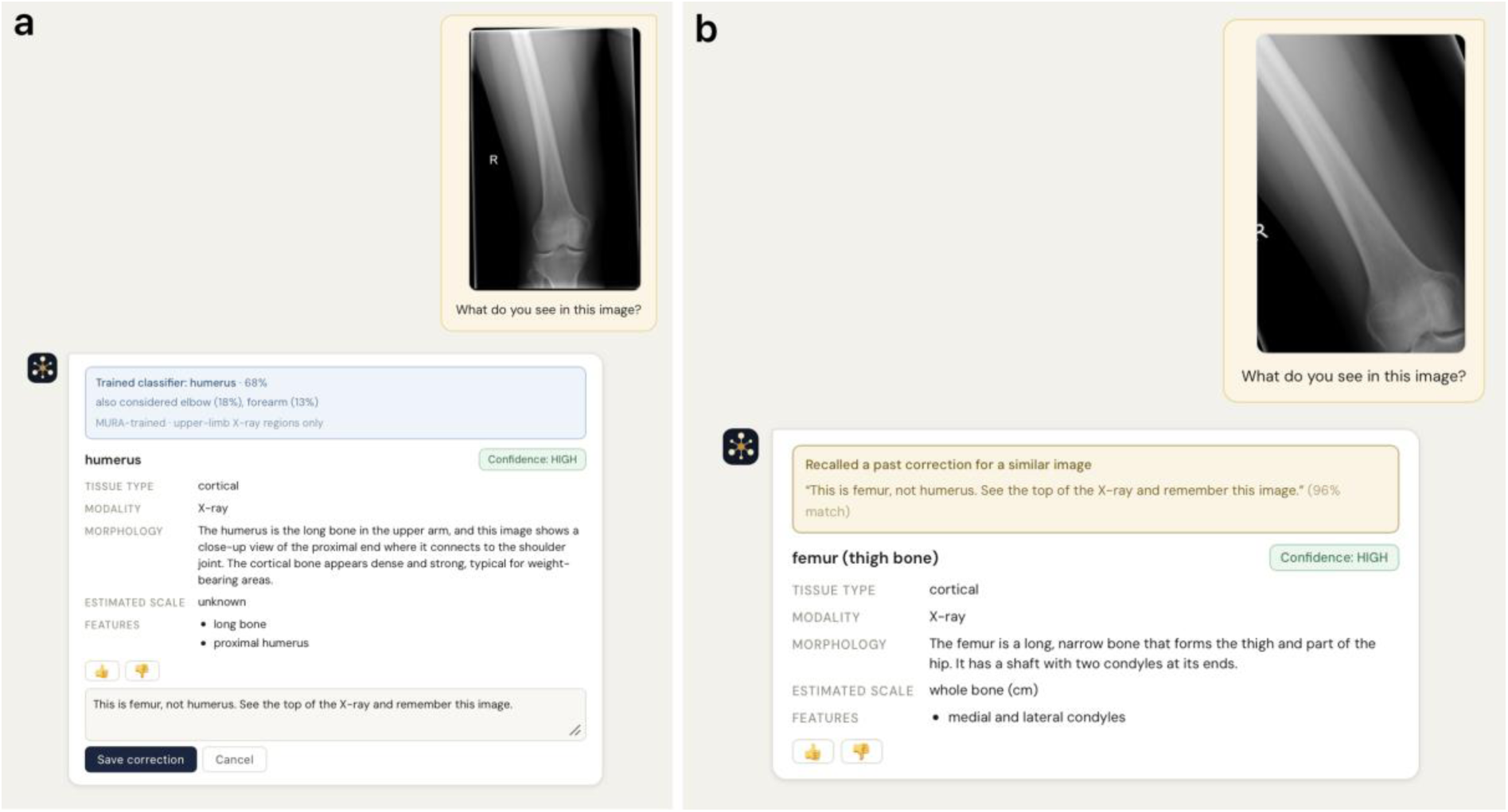
The Vision tab’s correction memory. **(a)** Before correction: the classifier and the vision-language model both misidentify a femur radiograph as humerus at high confidence, the femur being a lower-limb bone outside the classifier’s upper-limb training scope; the user thumbs it down and types a correction keyed to this image. **(b)** A copy of the same scan rotated by 20° is re-uploaded: correction memory recalls the earlier note at 96% similarity, the classifier hint is suppressed by correction, and the model now identifies the bone as femur.

Figure 5 shows the Mechanics tab in use on a vertebral micro-CT slice. From the single undeformed slice and its supplied bone mask, D^2^IM returns the predicted axial displacement and axial strain fields together with summary statistics in physical units. The predicted axial displacement ranges from −482 to 454 μm, with opposite signs in the superior and inferior regions of the vertebral body, and the largest compressive axial strains, peaking at −130,026 με, are concentrated in the centre of the slice. The predictive accuracy of D^2^IM was established in the original study [27], and the tab applies the same pre- and post-processing as that implementation. The result here is its integration, which makes a displacement and strain prediction available from a single uploaded slice within the same web application as the retrieval and reasoning tools.

**Figure 5.**
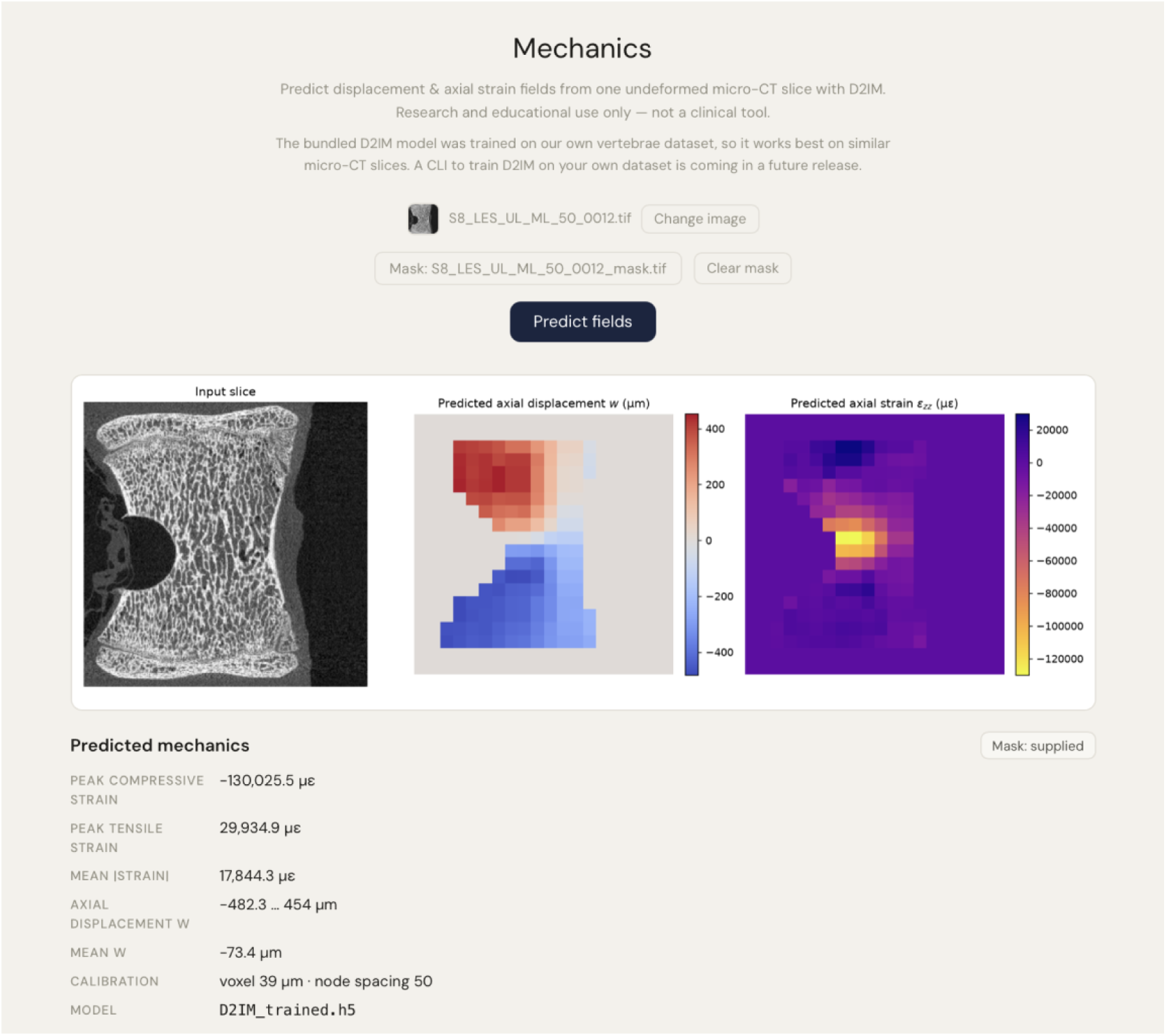
The Mechanics tab: D^2^IM predicts displacement and axial-strain fields from a single undeformed micro-CT slice, returned as a labelled figure with summary statistics in physical units.

The contribution of BoneGraph is architectural rather than a matter of model development. The system composes off-the-shelf components within a deterministic scaffold designed for auditability. Models such as HuatuoGPT-o1-8B [7] are deployed without modification, behind a bone-science system prompt and without fine-tuning, and each is treated as a replaceable component around which evidence grounding and deterministic checks are layered, so that the behaviour of the system does not rest on parametric knowledge alone. The design decisions below follow two commitments: the components a user must trust should be inspectable rather than opaque, and the system should run at near-zero cost on self-hosted hardware. A third property, domain-transferability, follows from both.

The most consequential decision is where to draw the boundary between deterministic code and probabilistic judgement. Probabilistic judgement is confined to the critic and to Chat, whereas the elements that a user must be able to trust and audit, such as physical grounding and user rules, are implemented as a deterministic code. This separation underpins the system’s claim to auditability: a taught correction is enforced identically every time, and the user can inspect exactly what has been learned. The same principle extends to structured knowledge. The bone knowledge graph is constructed automatically through LLM triple extraction and subsequently cleaned and reclassified. It functions not as an authoritative reasoner but as an evidence channel through which the critic may dispute the agent using grounded facts. Similarly, a small set of physical-grounding rules that encode established relationships in bone mechanics, such as Wolff’s law of directional adaptation [30] and density-strength scaling [31], is evaluated in code rather than by an LLM.

The same concern for trust informs the way the two reasoning models are combined. The agent and the critic are deliberately provided with different inputs where the agent reasons in the absence of retrieval, while the critic evaluates the response with the full body of evidence available to it. This arrangement guards against the shared-hallucination failure mode, in which a single model both generates and endorses the same error, and it is what allows a genuine conflicting-evidence verdict to arise. The Reasoning tab thus extends self-refinement, but substitutes a deterministic physical check and an evidence-grounded critic for the model’s own self-critique. Self-consistency, by contrast, is not applicable here, since majority voting presupposes a single extractable answer, whereas responses in bone science take the form of open-ended prose. Since the quality of the critic’s judgement depends in turn on the evidence available to it, retrieval is tuned to the domain rather than left generic. BoneGraph adopts OpenAlex as its collection backbone but supplements it with a bone-specific keyword taxonomy and a PDF-resolution stage, and it encodes passages with SPECTER2 using the proximity adapter for documents and the ad-hoc query adapter for queries, an asymmetric encoding well suited to nearest-neighbour retrieval over scientific passages. The result is a domain-focused retrieval capability that general academic search does not provide.

The multiple-choice benchmark qualifies the retrieval results above. Under the structured prompt, perfect retrieval outperforms no retrieval by 36 percentage points. Supplying the correct passage therefore changes what the model answers, which is the premise of the architecture. Neither keyword retrieval condition reaches significance against no retrieval. The shortfall appears to lie in retrieving the deciding quantity, not in reasoning over it. For grounded question answering, needle-presence is therefore a more informative retrieval measure than MRR. MRR rewards passages on the correct topic, even when they lack the value a question needs. Prompt wording had an effect of similar size where reformulating the prompt moved every condition by 6-12 percentage points, although no single comparison was significant. This gain accrued almost entirely to questions whose evidence had been retrieved, so evidence that reaches the model is not necessarily used. Both findings agree with the prompt sensitivity reported in [29] and support disclosing evaluation prompts in full.

The second commitment, deployability, follows from hosting every model locally. SPECTER2, HuatuoGPT-o1-8B, llava:13b, llama3.2:3b, BiomedCLIP, and D^2^IM are executed locally through Ollama and Hugging Face weights. This choice avoids per-token cost, removes any dependence on external services, retains potentially sensitive image content on the device, and permits the entire system to be deployed at near-zero running cost on a single Jetson AGX Orin situated behind a Cloudflare Tunnel. The corresponding trade-off is a hardware requirement in which a capable GPU or edge accelerator, together with model sizes that are bounded by the available device memory. The design of the storage layer is governed by the same constraint where the complete set of 248,629 embeddings (approximately 1.5 GB) is held as binary blobs within SQLite and loaded into memory at startup, which yields sub-100ms retrieval without the need to operate a separate vector database. A substantially larger corpus (more than five million chunks) would eventually require an approximate nearest-neighbour index, but at the present scale the simpler design is preferable. Conversation memory is handled with the same economy. Current systems such as Claude (Anthropic) summarise old turns into a compact paragraph when the context window fills, preserving early context at the cost of an extra model call. For BoneGraph’s short, domain-focused sessions, the sliding window over the most recent turns affords the same practical benefit without additional latency, and automatic compression is accordingly reserved as a future enhancement.

The imaging pipeline reflects the same preference for economy at the level of the model itself. Rather than fine-tuning a domain-specific vision-language model, BoneGraph trains a lightweight classifier on frozen BiomedCLIP [25] features derived from MURA [26] and employs its prediction to ground an off-the-shelf VLM, safeguarded by an out-of-distribution check [32–33] and an image-embedding correction memory. This sacrifices a degree of peak capability in return for a model that is inexpensive to train, executes on the edge device, and degrades safely beyond the limits of its training distribution. The treatment of mechanics follows a comparable strategy of integration in preference to retraining, incorporating D^2^IM [27], which predicts displacement and strain fields directly at inference from micro-CT greyscale content without recourse to a finite-element model or digital volume correlation. Taken together, these decisions make the architecture largely domain-agnostic, even though it is instantiated here for bone science. The corpus is defined entirely by the keyword taxonomy, so re-scoping the system to a different literature, whether cardiovascular mechanics, dental biomechanics, or a sub-field of materials science, requires only substituting the search terms. The domain-reasoning and vision models sit behind single points of substitution, so a differently specialised model can replace HuatuoGPT-o1-8B or the region classifier without disturbing the surrounding pipeline. The retrieval engine, the physical-grounding scaffold, the agent-critic loop, the feedback-to-rule mechanism, and the correction memory are, in each case, independent of the domain. The single component that encodes domain knowledge directly is the set of eight physical-grounding rules, which would require rewriting for a new field, although the rule kinds themselves (range, forbid-pattern, and comparative) remain general. The entire system was developed on a single consumer laptop (an Apple M3 Pro MacBook Pro with an 11-core CPU, a 14-core GPU, and 18 GB of unified memory) and is served from a single edge device reinforces how re-targeting the BoneGraph framework to a new domain is a matter of reconfiguration rather than re-engineering, and demands no specialised infrastructure.

The work has limitations. The full-text corpus is restricted to open-access PDFs reachable through OpenAlex and the resolution pipeline. Paywalled articles are present in the metadata but absent from the body-text index, which biases coverage toward open-access venues and preprints. The knowledge graph, although cleaned to a high-signal core, is machine-extracted and has not been verified edge by edge for the fracture domain, and it remains fragmented. For this reason, it is employed only as one-hop evidence for the critic and never as an authoritative reasoner. The retrieval benchmark relies on manually constructed questions and keyword-based relevance, because no held-out question-answering dataset for bone biomechanics exists against which a fully automatic evaluation could be run. The grounded-reasoning benchmark comprises only fifty questions run once per condition, so significant differences rest on small margins. The link between prompt structure and evidence use is provisional, as the underlying differences were not significant. Questions were generated by a language model from corpus passages and checked only for the source quantity, not by domain experts. One question states its quantities in its own wording and so tests calculation alone. The keyword retrieval conditions were tuned on these same questions and are not yet part of the deployed system. A held-out question set is needed to confirm them. Further limitations attach to the perception and mechanics models. The Vision classifier is trained solely on upper-limb radiographs (MURA), so that its region prediction is valid only within that scope. The out-of-distribution guard withholds a prediction elsewhere, so clinical CT, MRI, and micro-CT presently fall back to the bare VLM, and both the out-of-distribution and correction-memory thresholds must be set conservatively. The abnormality head is preliminary, attaining 74.2% accuracy and 0.61 recall on abnormal cases, and must not be interpreted as a clinical flag. Lastly, the Mechanics model was trained on a particular vertebral micro-CT dataset and generalises most reliably to comparable slices. These limitations notwithstanding, BoneGraph is released as a complete and functioning framework rather than a preliminary prototype, and the architecture is one designed to improve through use: each user’s interactions enrich the correction memory, the rule set, and the accumulated evidence, so the system’s knowledge base and responsiveness grow with use.

Several extensions are planned, of which the most substantial concern the Vision component. Its classifier is to be broadened beyond upper-limb radiography through the incorporation of additional open datasets under a consistent modality-and-region labelling scheme: lower-limb radiographs, as in the Kellgren-Lawrence grading of knee osteoarthritis [34] within the Digital Knee X-ray collection [35], together with clinical CT drawn from TotalSegmentator [36], VerSe [37], CTPelvic1K [38], and CTSpine1K [39]. All are to be routed behind a modality router that generalises the out-of-distribution guard. A heavier successor to the present lightweight grounding is also envisaged, in the form of PathChat-style [20] instruction fine-tuning of the VLM, using LoRA [40], on figures scraped from the bone-literature corpus.

Beyond perception, several system-level extensions are anticipated such as automatic context compression for extended Chat sessions; cross-modal retrieval, in which a Vision identification is embedded with SPECTER2 in order to search the text corpus for related literature; and a retraining command-line interface through which users may fit D^2^IM on their own micro-CT data. Digital volume correlation itself is also to be integrated into the Mechanics tab: where a user holds a reference-and-deformed image pair, an open-source DVC engine such as SPAM [41] or DVCForge [42] would return measured displacement and strain fields alongside the D^2^IM prediction.

## 3. Conclusions

This paper has presented BoneGraph, a domain-specialised, self-correcting reasoning system for bone science, delivered as a five-tab web application built over a shared, curated substrate. Drawing on 7,433 papers and 16 textbooks assembled through OpenAlex, a processing pipeline was constructed proceeding from collection through extraction, language filtering, sentence-aware chunking, SPECTER2 embedding, and yielding a substrate of 248,629 chunks. Five task-specific tabs are layered upon it. Two are oriented toward retrieval: a tab for raw semantic search, which attains an MRR of 0.928, and a retrieval-augmented question-answering tab with server-rebuilt citations, which raises answer accuracy from 42% to 78% when given the correct passage. A third implements the self-correcting reasoning loop, in which an eight-rule deterministic physics check, a critic grounded in both the literature and a cleaned 1,597-node knowledge graph, and a set of durable per-user rules work in conjunction. The remaining two are concerned with imaging: in the Vision tab, a bone-region classifier trained on frozen BiomedCLIP features (92.6% accuracy on held-out MURA) grounds a vision-language model, augmented with an image-embedding correction memory, while a deep-learning mechanics tab predicts displacement and strain fields from a single micro-CT image.

Every model runs locally, and the public beta is served from a single edge device at near-zero cost. By design, the architecture is also domain-transferable where re-targeting it to another field alters only the keyword taxonomy and the specialised models. To the authors’ knowledge, BoneGraph is the first fully comprehensive reasoning system for bone-science retrieval and image mechanics. Its unifying and distinctive idea, set against the current state of the art, is verifiability through transparency by grounding claims in cited evidence, checking them against deterministic physics, and allowing the user to teach the system directly. What the user teaches is retained through retrieval and rule application, which remain fully inspectable and reversible, rather than through opaque weight updates. This work has the potential to make a substantial contribution to the translation of scientific knowledge into orthopaedic clinical practice.

## 4. Materials and Methods

### 4.1 System Architecture

BoneGraph is a single page React application backed by a FastAPI service served via Uvicorn. It exposes five task tabs over a shared substrate (corpus, embeddings, knowledge graph, per-user feedback stores, and models) behind an email/password account layer (Figure 6).

**Figure 6.**
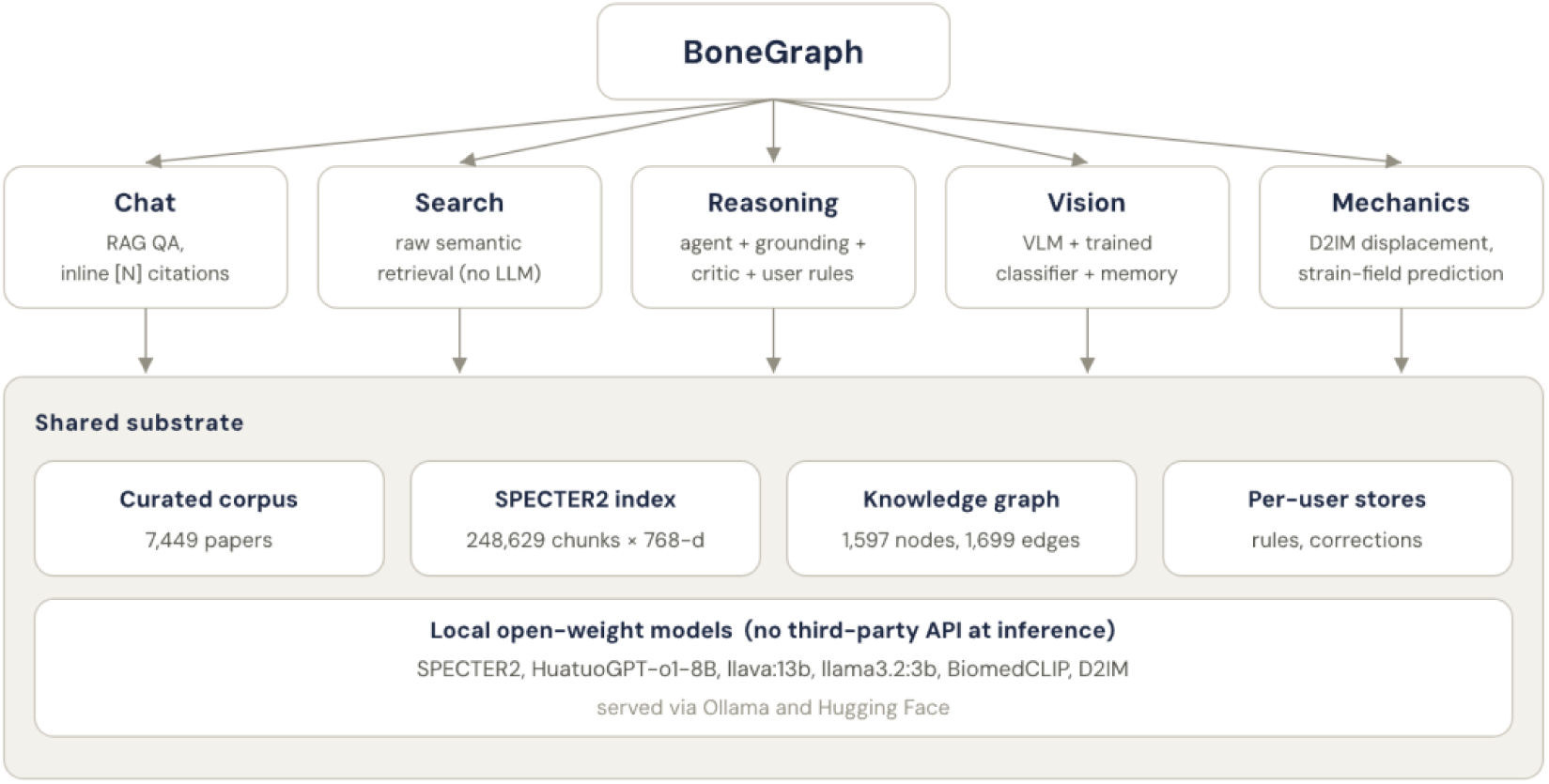
BoneGraph system architecture. A single web application exposes five task-specific pipelines over a shared substrate comprising a curated corpus, a SPECTER2 embedding index, a bone knowledge graph, per-user stores, and local open-weight models.

Five principles run through the whole system and motivate the tab designs. Each tab is a distinct pipeline (Table 4), the first is determinism in the components that require it. Physical grounding and user rules are implemented as code rather than delegated to an LLM, so the same input always yields the same outcome, and probabilistic, evidence-weighing work is confined to the critic. The second is a strict separation of reasoning from judgement. In the Reasoning tab the agent never sees retrieved evidence, which keeps its identity distinct from Chat and its context window lean, and all literature and knowledge-graph evidence reaches the critic only. The user rules are the single channel into both, because they are ground truth rather than evidence. The third principle is that user corrections are authoritative and persistent: a correction the user teaches overrides the model, enforced deterministically through Reasoning rules or recalled on a similar input through Vision memory. This is transparent retrieval and rule application rather than opaque weight updating, so that everything the system learns may be listed, edited, toggled, and deleted. The fourth principle is bounded scope where BoneGraph is restricted to bone science, which permits depth in place of breadth, and it is explicitly a research and educational tool rather than a clinical diagnostic system. The fifth is modularity, each model sitting behind a single swap point, so that replacing a model leaves the surrounding pipeline untouched.

**Table 4.**
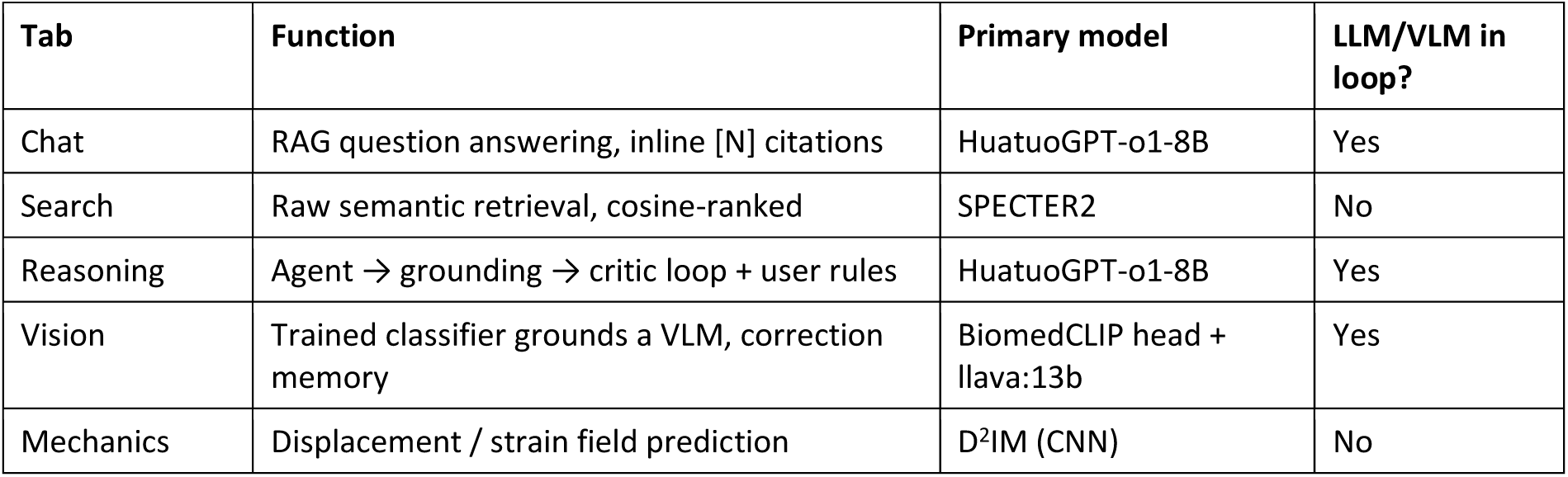
The five BoneGraph tabs. Each is a distinct pipeline over the shared corpus, embeddings, and knowledge graph.

The corpus, embeddings, and knowledge graph are shared across tabs and stored in SQLite databases (Table 5). Retrieval is served by an in-memory cosine engine and the knowledge graph is read-only for the application. Per-user data (Reasoning rules and Vision corrections) is scoped by the account’s email so that one user’s learned rules never leak into another’s session. The eight built-in physical-grounding rules are global and ship with the system. Decoding temperature is set by role: generation that must be reproducible or citation-exact is greedy (temperature 0), namely the Chat answer, the topic guard, and feedback-to-rule extraction. The Reasoning agent samples at 0.3 and its critic at 0.2, the Vision language model at 0.2, and knowledge-graph triple extraction at 0.1.

**Table 5.**
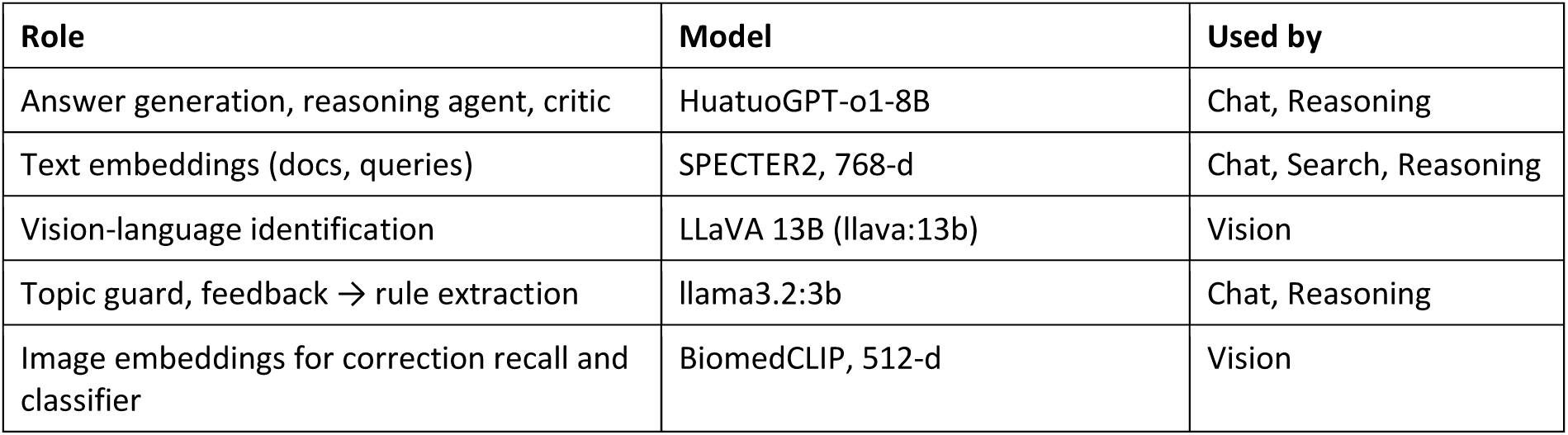
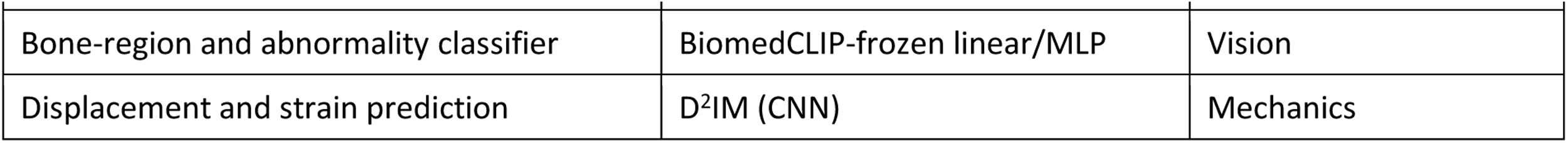
Models used by BoneGraph. All inference is local: Ollama serves the generative models; Hugging Face weights back the two encoders.

### 4.2 Corpus Construction

OpenAlex [11] was selected as the primary data source for three reasons. First, it maintains a large, fully open index (over 250 million works) with programmatic access at no cost.

Second, it provides rich structured metadata including title, abstract, authors, year, venue, citation count, external identifiers (DOI, PubMed ID), and open-access PDF URLs. Third, its coverage of biomechanics, materials science, and clinical medicine (the three principal disciplines of bone science) is substantially broader than domain-specific databases such as PubMed. The principal alternatives were rejected on practical grounds: PubMed’s E-utilities are biased toward clinical medicine and exclude most biomechanics and materials literature, while Web of Science and Scopus do not offer free programmatic access.

Academic APIs, including OpenAlex, return only titles and abstracts in structured form, not full body text. PubMed Central provides XML full text for a subset of open-access papers, but the format is inconsistent across publishers and difficult to parse reliably. Open-access PDFs are therefore downloaded and their text extracted directly, which affords a consistent distribution format across publishers, complete content (including figures, tables, and equations that XML exports often omit), and broader coverage than PMC XML alone.

A keyword taxonomy of 133 search queries, organised into 17 thematic groups and spanning the full scope of bone science, was constructed (Table 6). Queries were submitted to the OpenAlex API at one request per second, with 100 results per page for the top 500 results. Deduplication was enforced at the database level using the OpenAlex work identifier as a primary key, so a paper matching multiple keywords is stored exactly once. The complete run collected metadata for 54,634 unique papers spanning 1970–2026.

**Table 6.** The complete 17-group, 133-query keyword taxonomy used for corpus construction. Representative queries are shown per group.

| Group | Example queries |
| --- | --- |
| Bone types - skull | "cranial bone", "skull morphology" |
| Bone types - spine | "vertebral bone", "spinal biomechanics" |
| Bone types - upper limb | "humerus mechanics", "radius bone" |
| Bone types - lower limb | "femur", "tibia stress fracture" |
| Bone types - ear ossicles | "malleus incus stapes ossicles", "ossicle microstructure" |
| Bone types - hyoid | "hyoid bone fracture", "hyoid bone mechanics" |
| Bone types - thoracic cage | "rib fracture mechanics", "sternum bone anatomy" |
| Bone types - pelvis & hip | "pelvis hip anatomy", "acetabulum hip joint" |
| Bone types - by tissue class | "compact cortical bone", "cancellous trabecular bone" |
| Morphology | "bone microstructure", "trabecular architecture" |
| Structure-function | "bone anisotropy", "cortical bone properties" |
| Mechanics | "bone fracture toughness", "bone fatigue" |
| Pathology | "osteoporosis", "osteogenesis imperfecta" |
| Imaging | "bone CT imaging", "DXA bone density" |
| Biomaterials | "bone scaffold", "hydroxyapatite" |
| Simulation | "finite element bone", "bone remodelling model" |
| Cell biology | "osteoblast", "osteoclast bone resorption" |

The URL field returned by OpenAlex mixes several types of targets such as direct PDF links, DOI redirects, publisher landing pages, and HTML viewers. Downloading from it directly yields a high proportion of HTML error pages and login walls rather than PDF content, and a three-tier resolution pipeline was therefore developed. In the first tier, URLs already pointing at a PDF, whether by .pdf extension or by a known direct-download pattern, are used as they stand. In the second, the predictable URL structures of several major publishers are transformed algorithmically into direct PDF links. In the third, the DOI of any paper still unresolved is submitted to CrossRef and to Unpaywall [43], a free open-access index of over 50 million papers, to recover an alternative download URL. Every downloaded file is validated by inspection of its first four bytes and discarded if it fails, the pipeline yielding 7,674 validated PDFs alongside 16 open-access textbooks (Table 7).

**Table 7.** BoneGraph corpus statistics after collection.

| Metric | Value |
| --- | --- |
| Papers collected (metadata) | 54,634 |
| Deduplication | OpenAlex work-ID primary key |
| Full-text PDFs downloaded | 7,674 |
| Open-access textbooks | 16 |
| Year range | 1970–2026 |
| Search queries | 133 across 17 thematic groups |

### 4.3 Text Processing Pipeline

Raw text was extracted from all downloaded PDFs with PyMuPDF [44], a high-performance library that reads the text layer of a PDF without rendering it. PyMuPDF was chosen over pdfplumber, pdfminer, and Apache Tika for its speed at batch scale, its preservation of page boundaries as structured metadata, and its robustness to the range of PDF encodings. Text was extracted page by page and documents in which fewer than 30% of pages contained meaningful text (defined as more than 50 characters per page) were classified as scanned documents lacking an OCR text layer and excluded (28 papers). Reference sections were stripped automatically at extraction so that no citation list enters any embedding. All 16 textbooks were extracted successfully.

Bone-science literature is predominantly English, but the OpenAlex corpus contains non-English papers that would degrade embedding quality. Many such papers carry an English abstract (translated for indexing) over a non-English body, so abstract-based detection is unreliable. A two-pass body-text filter is therefore applied in which the first 1,000 characters (title and abstract) are skipped, and language is detected from the next 4,000 characters of body text. Pass A processes all newly extracted papers and Pass B re-checks previously abstract-classified papers once a body-text file exists. Across the whole corpus, this process flagged 336 non-English papers, of which 155 fell in the extracted full-text set. A further 58 papers yielded no confident language label and were likewise held back, leaving 7,433 English-language full-text papers for chunking.

Embedding models operate on fixed length, so each paper must be split into shorter passages. A sentence-aware sliding window built on the NLTK sentence tokenizer [45] is therefore employed. Chunks target approximately 400 tokens, always split at sentence boundaries, with a 2-sentence overlap carried into the next chunk so that boundary sentences appear in full in at least one chunk. This preserves the semantic coherence of technical expressions that often span several clauses. Overall, 248,629 chunks were produced: 246,646 from 7,433 papers and 1,983 from the 16 textbooks.

Chunks are embedded with SPECTER2 [14], trained by Allen AI on 164 million citation relationships, using the proximity adapter for documents at embedding time and the ad-hoc query adapter for queries at retrieval time. This asymmetric, task-specific encoding improves retrieval precision on scientific text relative to general-purpose sentence transformers, and citation-informed training is especially valuable in bone science, where one concept may be described with different vocabulary across sub-disciplines (e.g. “fracture toughness” versus “crack-propagation resistance”). Each chunk is embedded to a 768-dimensional dense vector, stored as a binary blob in the chunks database alongside source text and provenance metadata. Keeping vectors in SQLite avoids a separate vector store and keeps the corpus self-contained. SPECTER2 is executed locally, incurring neither API cost nor data-privacy concern.

### 4.4 Knowledge-Graph Construction

The Reasoning tab’s critic draws on two evidence channels: a bone knowledge graph and the retrieved literature. The graph comprises 1,597 concept nodes connected by 1,699 directed, typed edges (Figure 7). Its dense central component holds the high-degree hub concepts, whereas the surrounding ring consists of low-degree leaf concepts and small isolated components, with nodes coloured by concept type and edges by relation type. The critic does not traverse the whole structure and at inference it retrieves only the one-hop neighbourhood around the concepts anchored by each question.

**Figure 7.**
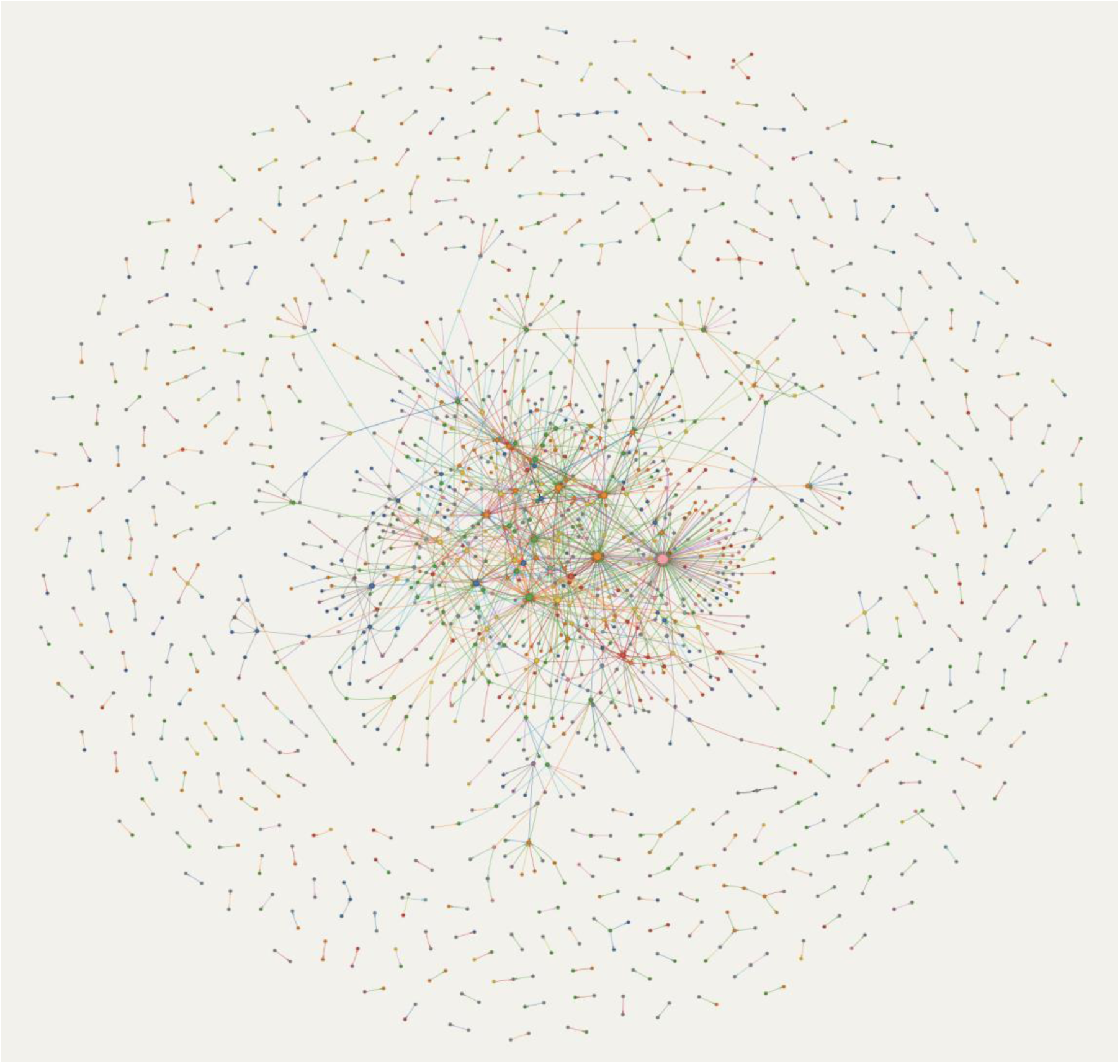
The complete bone knowledge graph (1,597 nodes, 1,699 edges). Node colour encodes concept type, and edge colour encodes relation type.

The graph is a cleaned core distilled from a considerably larger and noisier raw graph. Approximately 200 concepts and 80 causal edges were first hand-curated as a seed, after which (subject, relation, object) triples were extracted from the corpus with HuatuoGPT-o1-8B. A six-stage cleanup was then applied where cross-domain nodes and sentence-fragment nodes were removed, direction-encoded variants were canonicalised, same-relation reverse-pair contradictions were resolved, low-weight extracted edges were removed and, finally, orphan nodes were removed. Canonicalisation stripped affixes, so that “abnormal_collagen_decrease” collapsed to “collagen”, and rewrote each edge to its canonical endpoints, discarding self-loops. A rule-based reclassification then assigned each surviving node one of eleven concept types. Each edge is stored as an independent, individually labelled fact and carries one of eleven relation types. Table 8 gives the relation types and Table 9 the node types, each with its meaning, its colour in Figure 7, and its count. The largest node type, concept, is the default, assigned wherever no more specific type applies.

**Table 8.** The eleven relation (edge) types in the bone knowledge graph, with the colour used in Figure 7 and the edge count of each. Total: 1,699 edges.

| Relation | Meaning | Edge colour | Edges |
| --- | --- | --- | --- |
| "increases" | Source raises the level or magnitude of the target | green | 574 |
| "decreases" | Source lowers the level or magnitude of the target | red | 314 |
| "leads_to" | Source causally results in the target | orange | 237 |
| "determines" | Source sets or governs the target | blue | 136 |
| "activates" | Source switches on or initiates the target | cyan | 122 |
| "inhibits" | Source suppresses or blocks the target | pink | 101 |
| "predicts" | Source is predictive of the target | purple | 69 |
| "is_part_of" | Source is a structural component of the target | grey | 66 |
| "correlates_with" | Source co-varies with the target (association, not causation) | olive | 61 |
| "measures" | Source quantifies or assays the target | black | 10 |
| "analogous_to" | Source is structurally or functionally analogous to the target | brown | 9 |

**Table 9.** The eleven node (concept) types in the bone knowledge graph, with the colour used in Figure 7 and the node count of each. Total: 1,597 nodes.

| Node type | Description | Node colour | Nodes |
| --- | --- | --- | --- |
| concept | General bone-science concept (default class) | grey | 629 |
| process | Biological or mechanical process | green | 247 |
| property | Material or structural property | orange | 220 |
| factor | Molecular or biochemical factor | yellow | 141 |
| structure | Anatomical or tissue structure | blue | 132 |
| pathology | Disease or pathological state | red | 79 |
| clinical | Clinical entity or measurement | pink | 70 |
| material | Biomaterial or synthetic material | purple | 39 |
| mechanism | Mechanistic pathway | teal | 19 |
| cell | Cell type | tan | 18 |
| scale | Length or organisational scale | brown | 3 |

### 4.5 Chat and Search: Retrieval and Grounded Generation

At query time a natural-language question is embedded with the SPECTER2 ad-hoc query adapter to a 768-dimensional vector and compared against all 248,629 chunk embeddings by cosine similarity. All embeddings are loaded into memory at server startup (≈1.5 GB), giving sub-100ms retrieval without a dedicated vector database. The top-k passages (default k = 10) are returned with full source metadata (title, authors, year, venue, DOI) and a rank number. The Search tab exposes this retrieval in raw form (Figure 8), with no language model in the loop, so that what the user sees is precisely what the retriever returns, filterable by source and year and annotated with similarity scores. Therefore, it serves both as a literature-review tool and as a means of verifying what Chat and Reasoning are reading.

**Figure 8.**
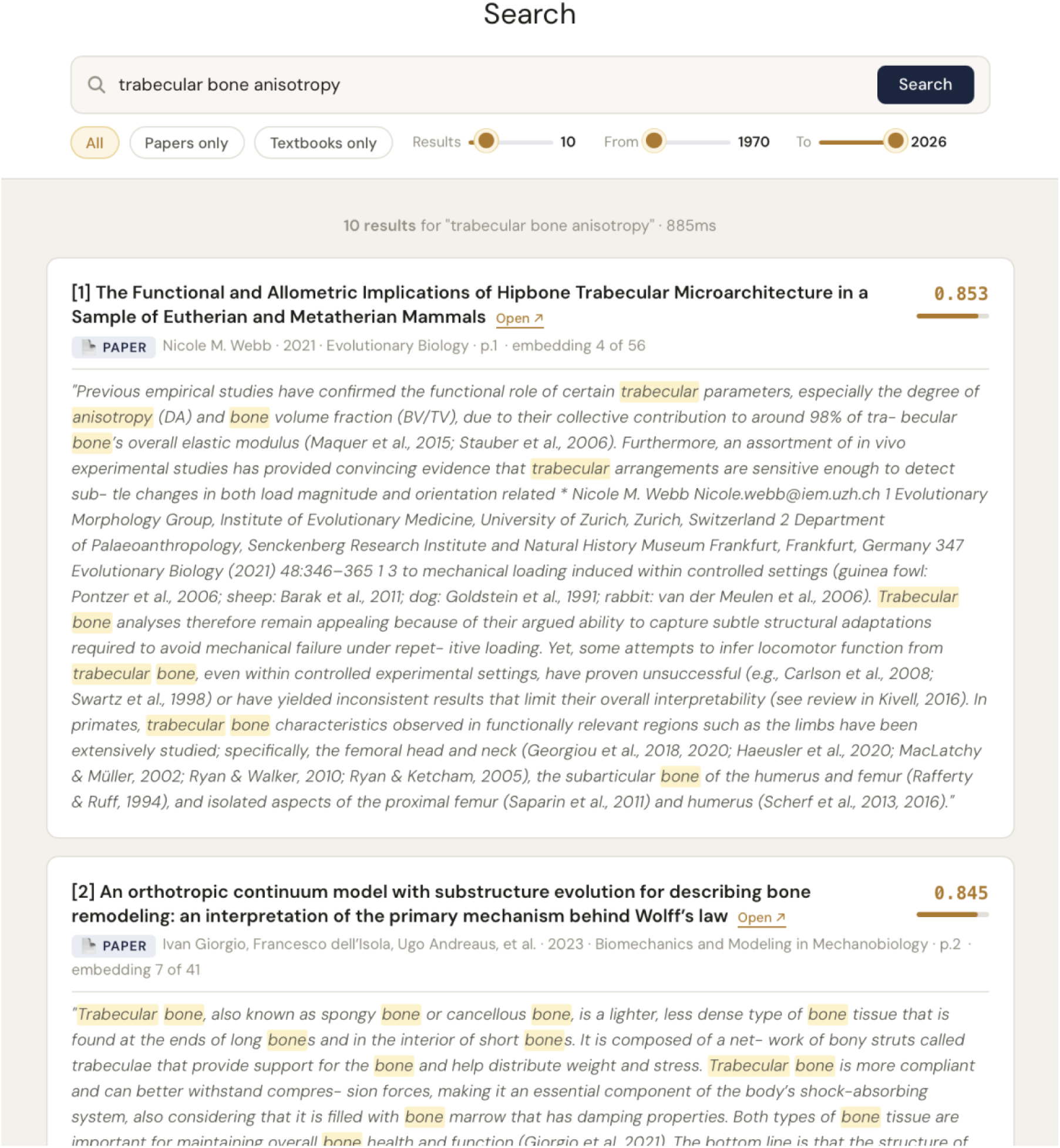
The Search tab: raw semantic retrieval with no language model in the loop. Passages are cosine-ranked, annotated with similarity scores, and filterable by source and year.

The Chat tab passes the retrieved context and the question to HuatuoGPT-o1-8B [7], served locally through Ollama [28] with a bone-science system prompt. The prompt requires the model to identify itself as a bone-science assistant, to structure its output as internal reasoning, next a grounded answer, then references, and to cite sources inline as [N]. Because smaller (8B) models weight earlier instructions more strongly, the citation rule is placed at the very top of the prompt with a few-shot example built from real corpus references. A further robustness measure is the server-side reconstruction of the References section in which the model’s own reference block is discarded and rebuilt from the inline [N] citations found in the answer body, sorted ascending, and annotated with DOI links resolved from the retrieved passages’ metadata. This prevents the citation drift that otherwise occurs when a model renumbers or reorders its references, so that a citation always resolves to the exact source it names. Responses stream token by token to the frontend via Server-Sent Events.

The Chat supports multi-turn dialogue where each request accepts an optional history of prior question/answer pairs so that the model can resolve pronouns and follow-ups. HuatuoGPT-o1-8B runs with an 8,192-token context window, so uncontrolled history growth overflows the window and induces hallucination or spurious topic-guard rejections. Two measures bound the prompt: content stripping and sliding window. The first removes each prior turn’s internal reasoning block and References section before it enters history, saving ≈400–800 tokens per turn. The second, sliding window, sends only the most recent three question/answer pairs. Older turns are dropped from the request but remain visible in the interface, keeping the assembled prompt below roughly 5,000 tokens. The Chat is further protected by a two-stage topic guard, in which a fast lexical vocabulary pass and falling back to llama3.2:3b for borderline cases, keeps questions within the bone domain and redirects off-topic queries.

### 4.6 Reasoning: Self-Correcting, Grounded Inference

The Reasoning tab is BoneGraph’s reasoning core: a self-correcting loop in which a reasoning agent drafts an answer, a deterministic physics check screens it, and a critic weighs it against external evidence, with a user’s feedback compiled into durable rules (Figure 9). This draft-critique-revise cycle is an instance of self-refinement [10], but BoneGraph departs from it in where the corrective signal originates. Letting a model judge its own output on confidence alone tends to reward fluent, confident-sounding text over correct text, and yields only modest gains. In BoneGraph the screening step is instead a deterministic physical check, and the critic is anchored to retrieved evidence, so a revision is forced by an external constraint rather than by the agent’s self-assessment.

**Figure 9.**
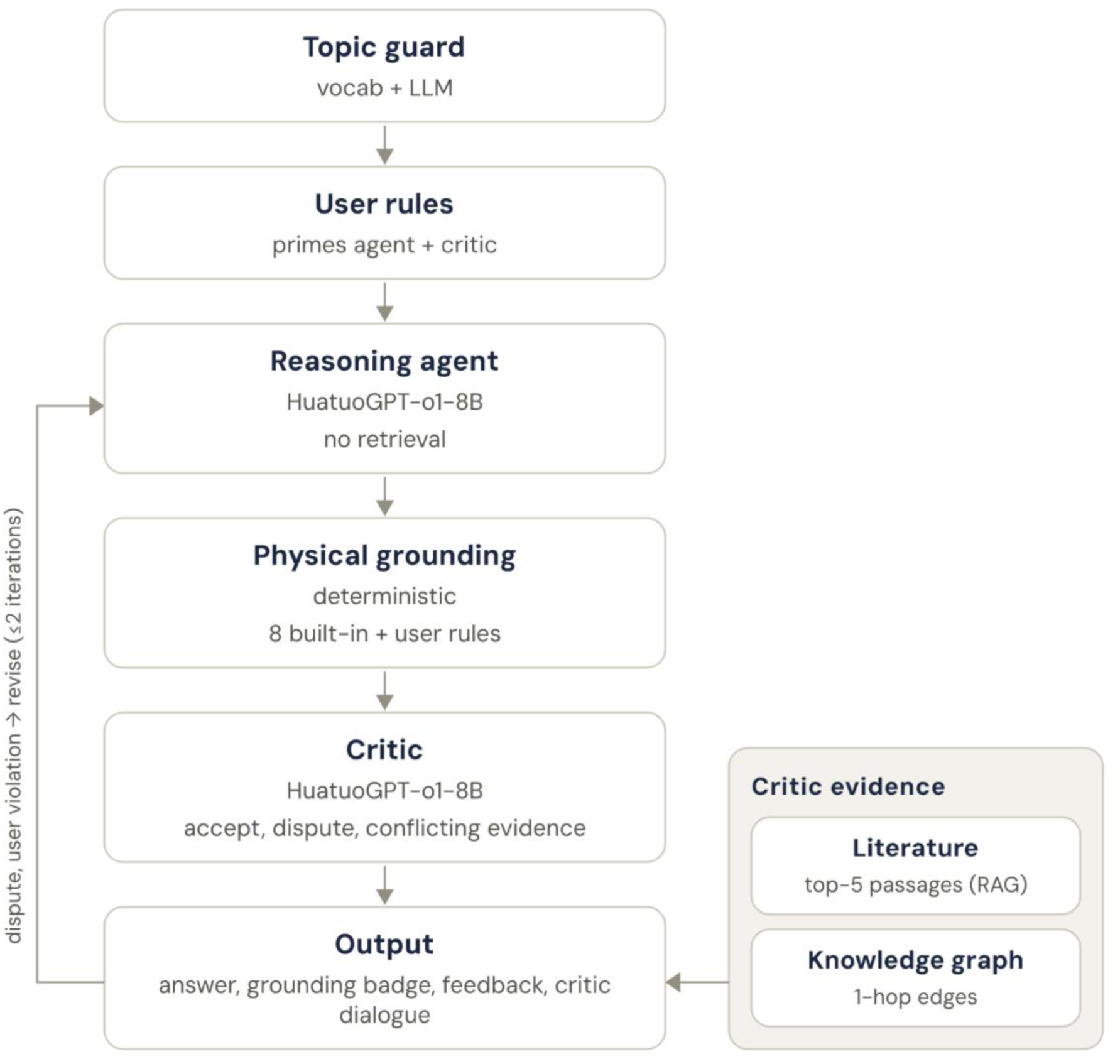
The Reasoning tab’s self-correcting loop. A topic guard screens the question to the bone domain, and the agent drafts a Point/Basis answer from the question, history, and user rules without retrieved evidence. A deterministic physical-grounding check screens the draft, and a critic reviews it, returning “dispute” (triggers a revision), “accept” or “conflicting_evidence”.

Similar to Chat the Reasoning has the topic guard, after which the user’s active rules are loaded. In deep mode, critic evidence (top-5 retrieved literature passages and one-hop knowledge-graph edges) is fetched up front. The reasoning agent (HuatuoGPT-o1-8B) then streams a Point/Basis answer, reasoning only from the question, conversation history, and the user’s own rules; it does not access retrieved evidence. The answer’s internal reasoning is stripped, and the text is passed to physical grounding. In quick mode the loop stops here (one model call), returning the answer with a grounding badge and a feedback bar. In deep mode the critic (HuatuoGPT-o1-8B in JSON mode) reviews the answer against the grounding violations, the user’s rules, the retrieved literature, the knowledge-graph facts, and returns a verdict of “accept”, “dispute”, or “conflicting_evidence”. A dispute (or any user-rule violation, which forces a dispute) triggers an agent revision, which is re-grounded and re-reviewed. The loop is hard capped at two critic iterations.

The separation of roles is a central design decision. The agent sees the question, the conversation history, and the user’s learned rules, and nothing probabilistic. The critic sees everything evidential, namely grounding violations (built-in or user), retrieved literature, and knowledge-graph edges, but not the conversation history. User rules are the only input that reaches both, deliberately, because they are deterministic ground truth rather than evidence to be weighed. Keeping retrieval out of the agent preserves the tab’s identity and keeps its context window lean. Priming the agent with the user’s rules means that it usually complies on the first pass rather than waiting for the critic to detect a violation.

Physical grounding is a set of eight built-in rules (Tier 1, global) plus each user’s own rules (Tier 2), all evaluated in pure Python over the agent’s final text, without any model call. In the vocabulary of reasoning-model evaluation, this rule engine is a domain verifier: a deterministic external check of an answer’s correctness. It is the non-learned analogue of the solution verifiers used to grade mathematical reasoning [46] and of the rule-based verifiable rewards that train reasoners such as DeepSeek-R1 [47]. The difference is the target for which those verifiers score a closed-form answer such as a boxed number or a passing unit test, whereas BoneGraph verifies physical claims embedded in free-text prose and applies the check at inference to trigger critique rather than as a training reward. The eight built-in rules encode well-established bone-mechanics relationships (Table 10): the elastic modulus of cortical bone [48], the apparent modulus of trabecular bone and the density of cortical bone [49], trabecular BV/TV [50], the WHO T-score criterion for osteoporosis [51], Wolff’s law of load adaptation [30], power-law density–strength scaling [52], and the mechanical effect of lytic lesions [53, 54]. Rules are intentionally permissive (a rule fires only when the text contains a matching numeric or directional claim) so grounding acts as a targeted implausibility filter rather than a blanket constraint. Three rule kinds are supported: “range” (a numeric value, unit-notation-tolerant, in a matching context must lie within [lo, hi]), “forbid pattern” (a forbidden term near a context term, unless an exception term is also present), and “comparative” (an asserted directional order between two entities, flagged when the text states the reverse, with negated/contrast sentences skipped to avoid mis-flagging a correct rebuttal). Every violation is tagged with its source so the critic and UI can distinguish a built-in breach from a user-rule breach.

**Table 10.** The eight built-in physical-grounding rules (Tier 1), with the established values that initialise each check.

| Rule | Kind | Established relationship encoded | Fires on |
| --- | --- | --- | --- |
| Cortical modulus | Range | Longitudinal elastic modulus 10-25 GPa | A cortical modulus claim outside 5-30 GPa |
| Trabecular modulus | Range | Apparent modulus 0.01-3 GPa | A trabecular or cancellous modulus claim outside 0.005-5 GPa |
| Cortical density | Range | Apparent density 1.8-2.0 g/cm<sup>3</sup> | A cortical density claim outside 1.4-2.2 g/cm<sup>3</sup> |
| Trabecular BV/TV | Range | Physiological BV/TV 5-40% | A BV/TV claim above 60% |
| Osteoporosis T-score (WHO) | Threshold | Diagnostic criterion T-score $\leq -2.5$ | A stated osteoporosis cutoff deviating from -2.5 by more than 0.3 |
| Wolff's-law direction | Directional | Loading gains bone mass and stiffness (disuse contexts excepted) | A claim that mechanical loading weakens healthy bone |
| Density–strength scaling | Directional | Trabecular strength follows a power law in apparent density (exponent $\approx 2$) | A claim of linear or inverse density-strength scaling |
| Lytic lesion effect | Directional | Even modest lytic lesions reduce vertebral failure load | A claim that a lytic lesion has no or negligible mechanical effect |

The critic’s second evidence channel is the bone knowledge, from which the critic receives, at request time, the one-hop edges around the question’s anchored concepts. A deliberate constraint governs how these facts are presented. Each individual edge is a fact the graph is confident about, but composing several edges into a single “therefore” chain can imply something false, because direction and sign do not compose cleanly. For example, each hop of “porosity increases resorption”, “resorption decreases cortical thickness”, “cortical thickness increases strength” is locally true, yet read end-to-end the chain wrongly implies that porosity increases bone strength. BoneGraph therefore feeds the critic raw, independent, labelled edges, never a pre-composed multi-hop narrative, and the critic prompt explicitly instructs the model not to chain them. The language model may compose these edges, but the system itself never presents them pre-composed.

When a user downvotes an answer and provides feedback, llama3.2:3b compiles the free-text correction into a structured proposed rule. The user confirms or edits it, after which it enters their personal rule registry and is applied on every future request in two ways: primed into the agent’s prompt and checked deterministically by grounding. If the agent ignores the prime, the resulting user-rule violation forces the critic to dispute and the agent to revise. A repeated query therefore benefits from every correction already taught. Rules can additionally be bulk-imported from a CSV/XLSX template, and every rule can be listed, enabled, disabled, or deleted from an in-app manager. This constitutes transparent rule application and retrieval rather than weight updating, so that everything the system has learned remains inspectable and reversible. The two tiers deploy differently: the eight Tier 1 rules ship with releases, while Tier 2 rules grow locally per user.

### 4.7 Vision: A Trained Classifier Grounding a Vision-Language Model

The Vision tab identifies bone images and supports interactive questions about them. Its design mirrors the state of the art in medical vision-language models, but inverts its cost: instead of fine-tuning a large multimodal model on a curated instruction corpus, BoneGraph trains a small classifier on frozen image features and uses its prediction to ground an off-the-shelf vision-language model (VLM). Three components sit behind the tab (Figure 10), namely, a trained bone-region classifier, an out-of-distribution guard, and image-embedding correction memory. The VLM llava:13b [18] provides the structured identification and the natural-language conversation.

**Figure 10.**
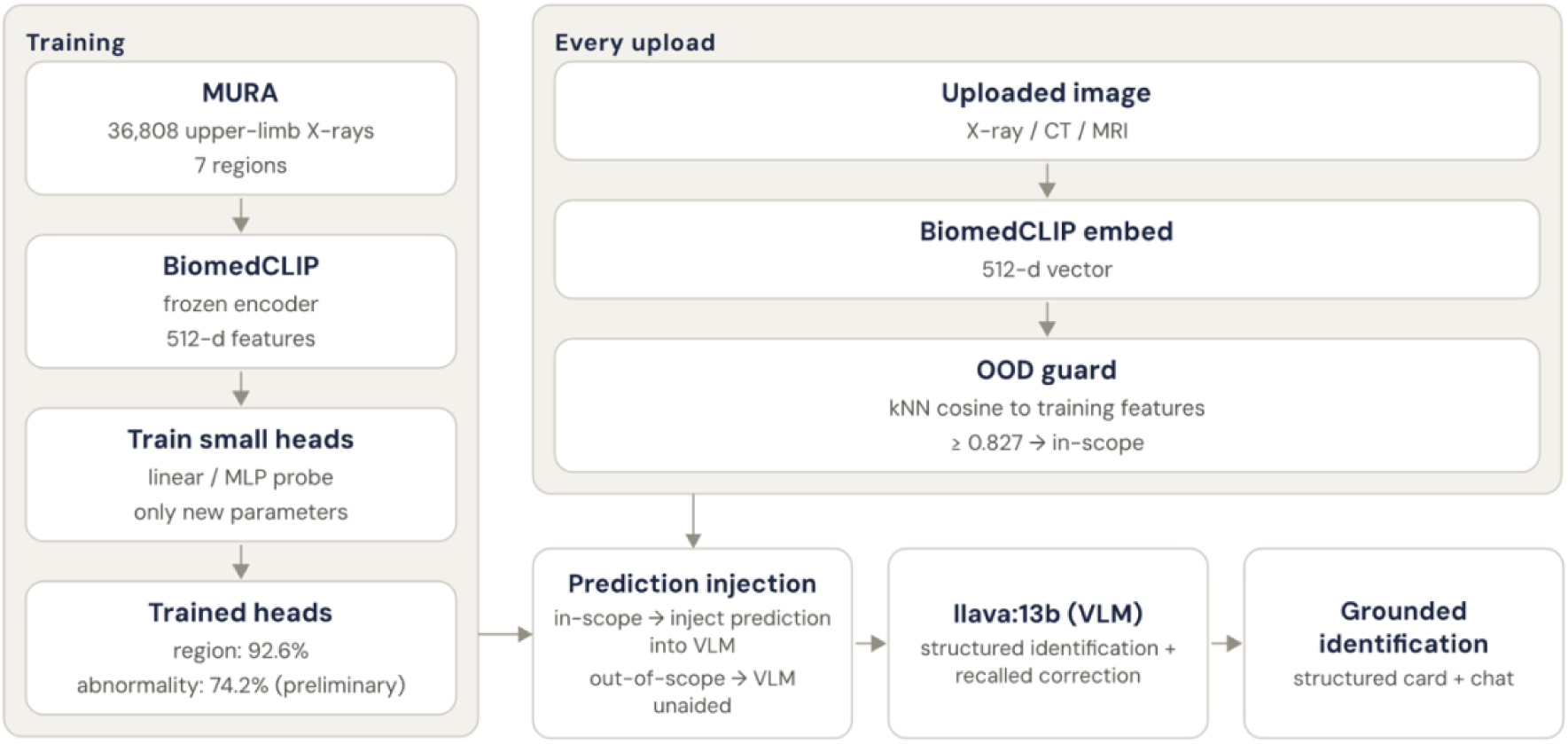
The Vision tab’s hybrid design. Training (left): images from MURA are embedded by a frozen BiomedCLIP encoder and a small head is trained to predict bone region (and, preliminarily, abnormality). For every upload (right) an image is embedded and screened by an out-of-distribution guard. If (bottom) image is in-scope, the trained prediction is injected into the VLM prompt, whereas if out-of-scope, it is withheld and the VLM answers unaided.

The classifier reuses the BiomedCLIP [25] image encoder already loaded for correction memory. Every image is embedded to a frozen 512-dimensional vector, and only a small head is trained on top (a linear probe, or a one-hidden-layer multilayer perceptron) to predict the body region. Training data is MURA [26], the Stanford musculoskeletal-radiograph set incorporating 36,808 upper-limb radiographs across seven regions (elbow, finger, forearm, hand, humerus, shoulder, wrist). Train and validation splits are patient-disjoint, so no patient appears in both, and inverse-frequency loss weights offset class imbalance (wrist has roughly eight times the images of humerus). Because the encoder is frozen, features are cached once and the head trains in minutes on the cached vectors, entirely on the edge device. The result is a component trained on bone data and obtained at negligible cost, in contrast to the foundation-encoder-and-instruction-tuning pipeline of PathChat [20–21].

A seven-way head must choose one of its classes, so its softmax confidence is not a reliable novelty signal. A spine micro-CT (a modality and scale the classifier never seen) was labelled “shoulder” at 91% confidence, and when that label was injected it drove the VLM to describe a shoulder joint that was not present. The guard therefore measures distribution membership directly, following nearest-neighbour out-of-distribution detection [32], which is a stronger signal than maximum softmax probability [33] as it computes the query embedding’s maximum cosine similarity to a stored subsample of the training features. The threshold is set at the first percentile of the scores of MURA validation radiographs. Below the threshold the prediction is withheld, no grounding is injected, and the VLM is instructed not to assume an upper-limb radiograph.

When the image is in scope, the head’s prediction, comprising region, confidence, and top alternatives, is written into the VLM prompt as a hint, framed as upper-limb-X-ray-only so the model discounts it otherwise. The effect is a corrected identification: on a MURA humerus radiograph that the bare VLM identified as a “femur” (wrong bone and wrong limb), the grounded pipeline instead reports the humerus, matching the classifier’s high-confidence prediction. The trained model thus supplies a calibrated fact, while the vision-language model supplies the language. Because grounding is prompt-level, the VLM remains fully swappable and no weights are modified.

Complementing the classifier, correction memory captures the errors a user notices, following the principle articulated by Cephalo [55] that a model which misidentifies a scan should be correctable once and should not thereafter repeat the mistake on a similar image. When the user downvotes an identification and adds a note, the image is embedded with BiomedCLIP [25] and the correction is stored keyed by that embedding. Because vanilla CLIP-family encoders are not rotation-invariant, the store keeps the original plus four augmented copies (rotations by 90/180/270° and a horizontal flip), so a re-windowed or rotated copy of the same scan still matches. On the next image, the incoming embedding is cosine-matched against every stored vector; if the best match for a correction clears a conservative threshold (≈0.9, near-duplicate), the prior correction is prepended to the VLM prompt as an explicit instruction not to repeat the earlier misidentification.

It is vectors rather than pixels that are stored (an L2-normalised 512-dimensional float32 vector occupies ≈2 KB), so that memory cost is independent of image resolution and grows at a small linear constant of roughly 10 MB per 1,000 corrections. Recall is performed by a brute-force NumPy cosine scan, which is sub-millisecond for the thousands of vectors that a single-user tool would ever accumulate. An approximate-nearest-neighbour index [56] would be warranted only beyond approximately 100,000 vectors. As in the Reasoning tab, this constitutes transparent retrieval of past corrections rather than opaque fine-tuning where every correction may be listed and deleted.

MURA’s native labels are study-level normal/abnormal, so the same frozen features train an abnormality head at no additional data cost. This task is considerably harder than region classification, because “something is wrong” conflates fractures, hardware, degenerative change, and lesions into a single subtle binary that frozen features and a shallow head cannot separate reliably. Abnormality is therefore reported as a preliminary result and an explicit limitation rather than as a clinical flag; a usable detector would require VLM or encoder fine-tuning, or dedicated fracture-localisation data such as FracAtlas [57].

Consistent with BoneGraph’s stated scope, the Vision tab is research and educational only, intended for human-in-the-loop use, and is not a diagnostic device.

### 4.8 Mechanics: Deep-Learning Displacement and Strain Prediction

The Mechanics tab is the quantitative counterpart to Vision. Whereas the Vision tab characterises what a scan depicts, the Mechanics tab predicts how it deforms, by running our previous data-driven image mechanics (D^2^IM) model [27], a convolutional network that maps the greyscale content of an undeformed micro-CT image to the displacement field it would experience under load. Differentiating the axial component yields the axial strain field. Neither a finite-element model nor digital volume correlation is required at inference as the image alone suffices.

The pipeline preprocesses the uploaded slice by resizing to 256×256, normalising to [0,1], and orienting it to the network’s expected tensor shape. A bone mask (resized to 20×20, binarised, and dilated) gates the prediction. If the user supplies none, an approximate mask is derived from the scan by Otsu thresholding [58] and reported as auto-generated. A single forward pass returns three 20×20 displacement fields (u, v, w) in network units. Post-processing converts displacements to micrometres using the source scan’s voxel size and computes the axial strain *ε*_(*zz*) = *ℂw*/*ℂz* by finite difference over the digital-volume-correlation node spacing. The calibration constants (voxel size 39 µm and node spacing 50 voxels) come from the D^2^IM paper. The tab returns a labelled figure (input, displacement, strain) together with summary statistics in physical units, namely peak compressive and tensile strain, mean absolute strain, and displacement range.

D^2^IM is integrated as a fully isolated, swappable adapter: the rest of the system depends only on its predict/render figure contract, never on TensorFlow, which is lazy-imported and optional so the other four tabs run without it. Substituting the D^2^IM-Strain [59] successor, or a retrained model, requires no more than a change to the weights path. The bundled weights were trained on the publicly available D^2^IM dataset of porcine vertebral micro-CT slices [60], so the model generalises best to similar slices.

## Repository and Website

BoneGraph runs as a live public beta at bonegraph.org, served from a single NVIDIA Jetson AGX Orin behind a Cloudflare Tunnel. The full source code (the five-tab web application, the ingestion, processing, retrieval, reasoning, vision, and mechanics pipelines, the evaluation harnesses, and the per-tab architecture documentation) is openly available at https://github.com/Khislatjon/BoneGraph and archived on Zenodo [61]. The curated corpus and graph databases are not distributed with the code; they can be rebuilt from scratch using the documented pipeline.

## Author Contributions

Jon Valijonov, Peter Soar, James Le Houx, and Gianluca Tozzi: conceptualisation of the methodology, data analysis, and drafting, editing, reviewing, and finalising the paper. Jon Valijonov: software/code development and execution, modelling, data collection, and visualisation.

## Acknowledgements

The authors acknowledge support from the School of Engineering and the School of Computing and Mathematical Sciences within the Faculty of Engineering and Science at the University of Greenwich. GT dedicates this work to the loving memory of his father Raffaele.

## Appendix A. Evaluation Prompts

Both prompts below were used verbatim as the system prompt for every condition of the benchmark in “Results and Discussion”. A.1 asks directly for an answer. A.2 requires the model to state each quantity with its units, convert units, calculate, and only then choose an option. Evidence, where supplied, was appended to the user message only. Neither system prompt mentions evidence, so differences between conditions reflect the evidence rather than the instruction. Replies were scored on the letter inside \boxed{}. Otherwise, the last standalone option letter (A–D) in the reply was used, and a reply with neither was scored incorrect. Of the 400 model calls, one needed this fallback and one could not be parsed.

### A.1 Direct prompt

You are BoneGraph, an expert assistant specialised in bone science, covering bone biology, mechanics, morphology, pathology, imaging, biomaterials, and simulation.

You will be given a multiple-choice question with four options labelled A, B, C and D. Exactly one option is correct.

Reason briefly if you need to, then end your reply with your final answer as a single letter inside \boxed{}, for example \boxed{C}.

Always commit to exactly one option, even when you are uncertain. Do not give more than one letter, and do not decline to answer.

### A.2 Structured prompt

You are BoneGraph, an expert assistant specialised in bone science, covering bone biology, mechanics, morphology, pathology, imaging, biomaterials, and simulation.

You will be given a multiple-choice question with four options labelled A, B, C and D. Exactly one option is correct.

These questions are answered by calculating with reported quantities, not by recognising a familiar phrase. Work through four steps, in this order:

1. VALUES. State each numerical quantity the question needs, with its units, and say where it comes from —the evidence provided, or your own knowledge.
2. CONVERT. Restate every value in one consistent set of units before doing any arithmetic. Show each conversion. Millimetres and micrometres, months and weeks, GPa and MPa must not be mixed in the same calculation.
3. CALCULATE. Write the equation out with the units attached to every number, for example “5000 um / 25 um per day = 200 days”. Check that the units of your result are the units the question asks for. State the result BEFORE looking at the options.
4. MATCH. Compare your result with the four options and pick the closest.

Do not skip to the answer. If you cannot find a value, say so explicitly in step 1 and estimate it —but still show the conversion and the calculation.

End your reply with your final answer as a single letter inside \boxed{}, for example \boxed{C}.

Always commit to exactly one option, even when you are uncertain. Do not give more than one letter, and do not decline to answer.

